# Performance assessment of total RNA sequencing of human biofluids and extracellular vesicles

**DOI:** 10.1101/701524

**Authors:** Celine Everaert, Hetty Helsmoortel, Anneleen Decock, Eva Hulstaert, Ruben Van Paemel, Kimberly Verniers, Justine Nuytens, Jasper Anckaert, Nele Nijs, Joeri Tulkens, Bert Dhondt, An Hendrix, Pieter Mestdagh, Jo Vandesompele

## Abstract

RNA profiling has emerged as a powerful tool to investigate the biomarker potential of human biofluids. However, despite enormous interest in extracellular nucleic acids, RNA sequencing methods to quantify the total RNA content outside cells are rare. Here, we evaluate the performance of the SMARTer Stranded Total RNA-Seq method in human platelet-rich plasma, platelet-free plasma, urine, conditioned medium, and extracellular vesicles (EVs) from these biofluids. We found the method to be accurate, precise, compatible with low-input volumes and able to quantify a few thousand genes. We picked up distinct classes of RNA molecules, including mRNA, lncRNA, circRNA, miscRNA and pseudogenes. Notably, the read distribution and gene content drastically differ among biofluids. In conclusion, we are the first to show that the SMARTer method can be used for unbiased unraveling of the complete transcriptome of a wide range of biofluids and their extracellular vesicles.

## 1. Introduction

All human biofluids contain a multitude of extracellular nucleic acids, harboring a wealth of information about health and disease status. In addition to established non-invasive prenatal testing of fetal nucleic acids in maternal plasma^1^, liquid biopsies have emerged as a novel powerful tool in the battle against cancer^2^. Although in the past most attention was given to circulating DNA, its more dynamic derivate extracellular RNA may provide additional layers of information. However, RNA sequencing in biofluids is technically challenging. Low input amounts, large dynamic range, and (partial) degradation of RNA hamper straightforward quantification. While sequencing of small RNAs^3^ and targeted or capture sequencing of longer RNAs^4^ proved to be successful, studies using total RNA sequencing on biofluids are rare. To date, only a few whole transcriptome profiling attempts were made on urine, plasma or extracellular vesicles^5–9^, quantifying both polyadenylated and non-polyadenylated RNA transcripts. However, all these methods suffer from one or more limitations such as short fragment length, low amount of quantified genes or ribosomal RNA contamination.

The advantages of total RNA sequencing are plentiful. Indeed, detection is not limited to a set of pre-defined targets, nor to (3’ ends of) polyadenylated RNAs. Next to polyadenylated mRNAs, various other RNA biotypes including circular RNAs, histone RNAs, and a sizable fraction of long non-coding RNAs can be distinguished. In addition, the study of posttranscriptional regulation is possible by comparing exonic and intronic reads^10^. Altogether, this generates a much more comprehensive view of the transcriptome.

Here we aimed to assess the performance of a strand-specific total RNA library preparation method for different types of biofluids and derived extracellular vesicles (EVs). We applied the method on platelet-rich plasma, platelet-free plasma, urine and conditioned medium from human healthy donors, cancer patients or cancer cells grown *in vitro*. More specifically, the SMARTer Stranded Total RNA-Seq Kit – Pico Input Mammalian, including a ribosomal RNA depletion step at the cDNA level, was extensively evaluated. We found the method to be accurate and precise. Low-input volumes are technically feasible and the method allows the detection of several thousand genes of different classes.

## 2. Results

### 2.1. Read distribution drastically differs among biofluids

In a first experiment (Fig 1A), we sequenced platelet-rich plasma (PRP) and platelet-free plasma (PFP) from two different healthy donors. We collected blood in EDTA tubes, hence the ‘e’ in front of ePRP and ePFP throughout the manuscript. From each plasma fraction, two technical RNA extraction replicates were performed, resulting in four sequenced samples per donor. Because of the low input, between 53.0% and 88.2% of the reads were PCR duplicates (SupFig1). PCR duplicates arise when multiple PCR products from the same original template molecule bind to the sequencing flow cell. For better quantitative accuracy, we removed the duplicates for further analysis. The variation in PCR duplicate levels between plasma fractions is related to the amount and quality of input RNA. As we will illustrate below, ePRP has a higher RNA input concentration, which explains the lower number of duplicate reads compared to ePFP. After duplicate removal we mapped the remaining (deduplicated) reads to the reference genome (Fig 2A). Four categories of reads can be distinguished here: uniquely mapping reads, multi-mapped reads aligning to several genomic positions, reads that are too short to map, and unmapped reads. The number of unmapped and multi-mapped reads was similar between plasma with and without platelets. However, ePFP samples contain much more reads that are too short to map. As a consequence, ePRP contains approximately twice as many uniquely mapped reads, possibly the result of more intact RNA in platelets. However, when only considering these unique reads, more than 75% of them derived from mitochondrial RNA (mtRNA) in ePRP (Fig 2B). In contrast, ePFP contains at least three times less mtRNA and considerably more reads mapping to nuclear DNA. Finally, also the distribution between exonic, intronic and intergenic reads differs between platelet-rich and platelet-free plasma (Fig 2C).

**Figure 1.**
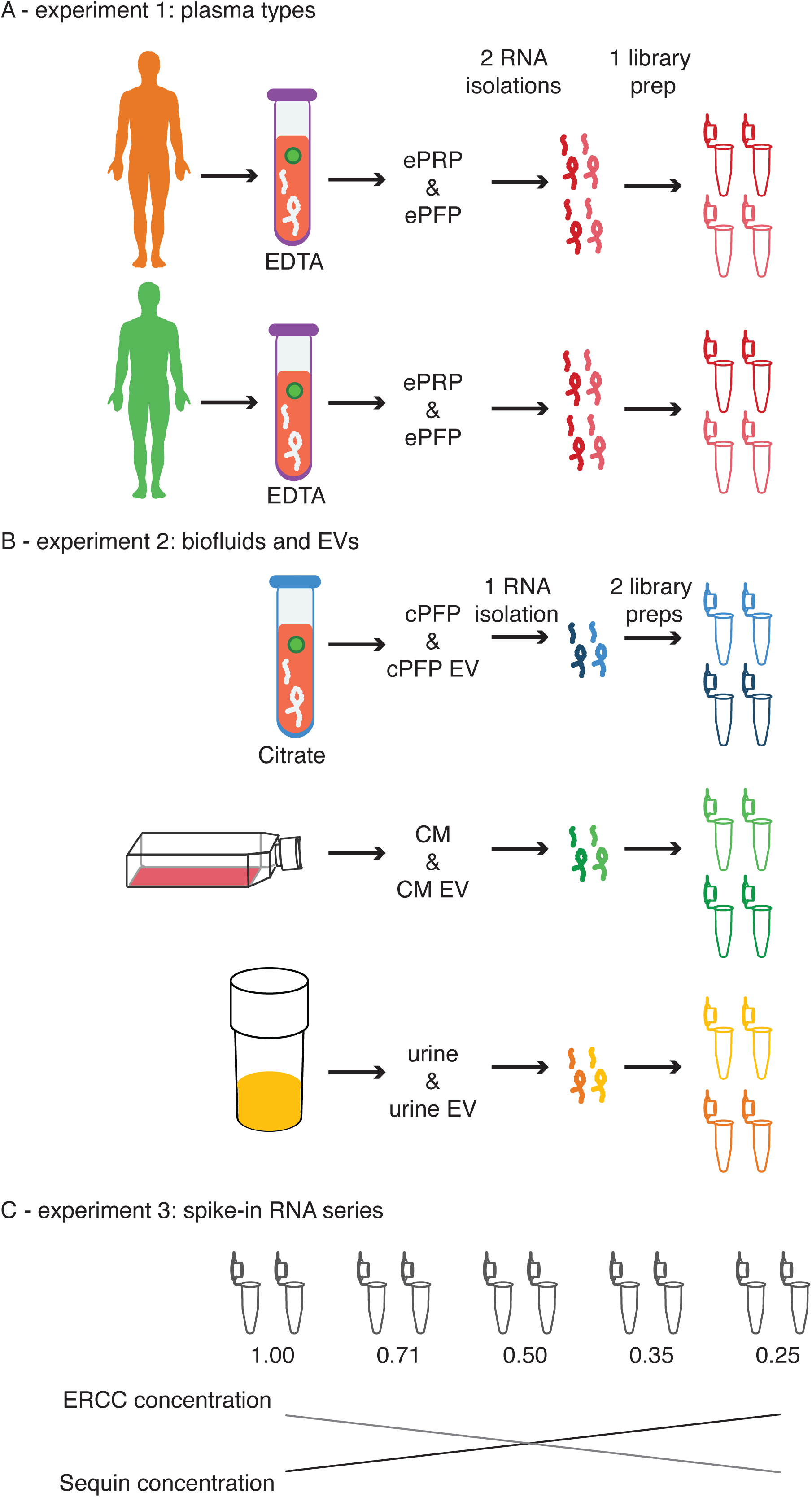
Schematic overview of the different experiments.

**Figure 2.**
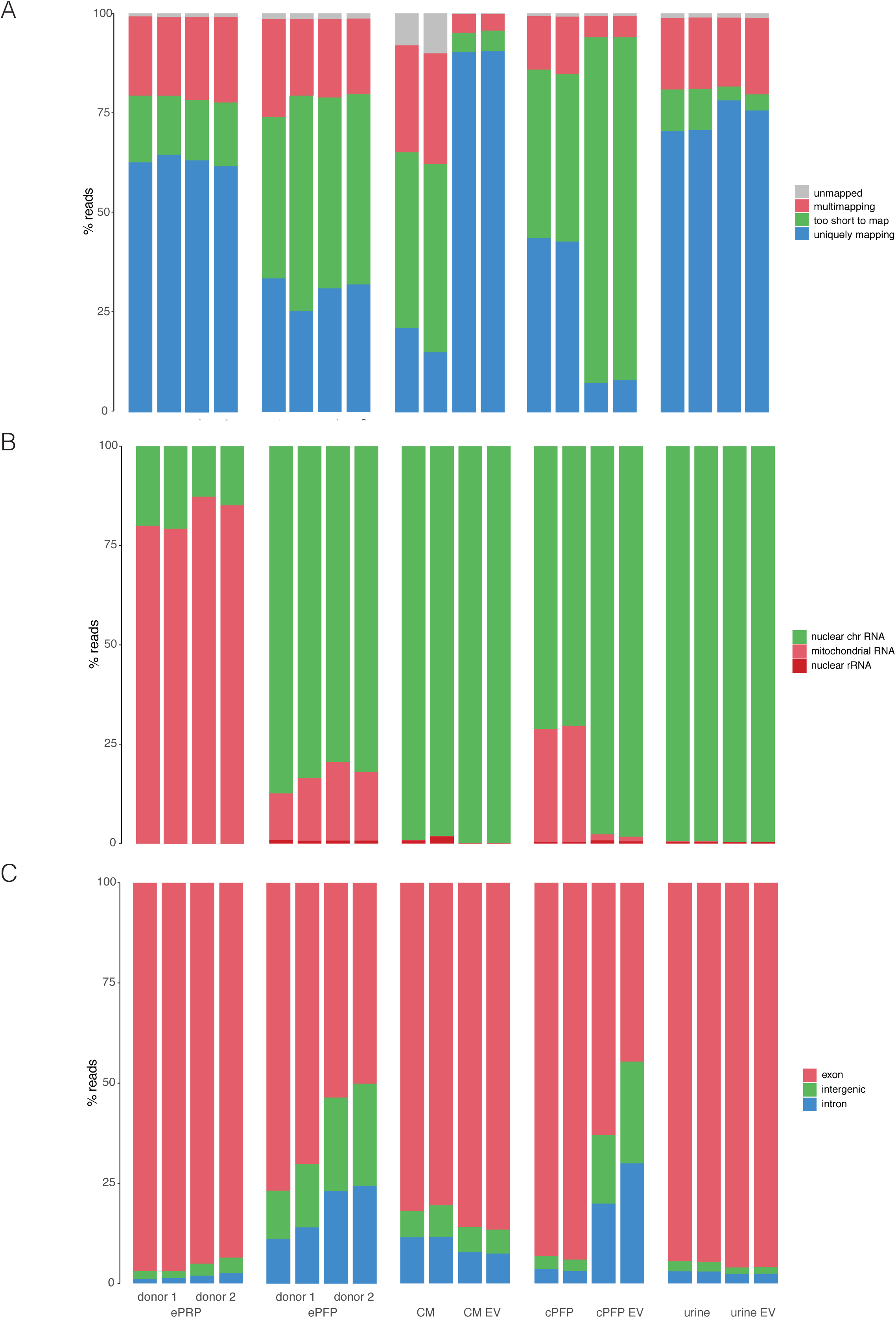
Read distribution of all libraries differs among samples. A) Percentage of reads assigned as too short to map, unique- or multi-mapping quantified with STAR. B) Percentage of reads derived from nuclear RNA, mitochondrial RNA and ribosomal RNA per sample quantified with STAR. B) Percentage of the reads originating from nuclear chromosomes derived from exonic, intronic and intergenic regions per sample quantified with STAR.

In the second experiment (Fig 1B), we sequenced conditioned medium from breast cancer cells (CM), platelet-free plasma from a third healthy donor collected in a citrate blood collection tube (cPFP) and urine from a prostate cancer patient. In addition, we purified EVs from these three fluids and performed extensive quality control using western blot, electron microscopy and nanoparticle tracking analysis (SupFig2). We sequenced the EV samples together with their fluids of origin. For this experiment, two technical replicates were introduced at the level of library preparation for each condition, resulting in 12 libraries. Because only one biological sample of each biofluid was included in this experiment, we should be cautious when generalizing differences among biofluids. With the exception of plasma, the number of PCR duplicates is lower in EVs compared to their parental biofluid (SupFig1). As mentioned earlier, the levels of PCR duplicates are typically lower in samples with higher input quality and concentration. But, as we will see in the next paragraph, RNA input amounts in EVs are not higher compared to their fluid of origin. Another explanation, at least partly, could be the protective effect lipid bilayers have on the quality of their RNA cargo. Interestingly, also mapping rates can differ substantially among biofluids and/or their EVs (Fig 2A). In our setup for instance, the fraction of unique reads ranges from 7.69% in cPFP EVs to 90.2% in EVs isolated from conditioned medium. When looking at the mapping properties of the unique reads, almost all samples mainly contain reads that map to nuclear DNA (Fig 2B). Only platelet-free plasma contains 25.8% mitochondrial RNA, comparable to the percentages that were generated in the healthy donors of the first experiment. Lastly, most reads mapping to nuclear DNA are exonic. The only exception here are cPFP EVs that contain a larger fraction of intronic and intergenic reads (Fig 2C). While the platelet-free plasma samples in the first and second experiment seem very similar, small differences may be introduced by blood collection tube (EDTA vs. citrate) and/or the use of distinct donors. Indeed, also in the first experiment the read distribution was to some extent donor dependent.

We subsequently investigated two other technical characteristics of our biofluid total RNA seq method: the level of strandedness and the inner distance between paired-end reads. In general, the method generates strand-specific sequencing reads in all the biofluids we assessed (SupFig3). The cDNA fragment sizes in the library range from 70 to 400 nucleotides, with a peak around 90 nucleotides for the plasma samples and around 180-190 nucleotides for the other samples. Notably, the plasma samples and derived EVs present with the shortest fragment length (SupFig4). In conclusion, we show for the first time that the SMARTer Stranded Total RNA-Seq method works in different human biofluids and their respective EVs. The method generates reproducible read distribution results for technical replicates, both at the RNA isolation and library preparation level. The results clearly differ according to biofluid sample type.

### 2.2. Spike-in RNA enables relative RNA quantification and fold change trueness assessment

In order to assess the quantitative aspect of the total RNA sequencing method, we added an ERCC RNA spike-in mix to all RNA samples prior to library preparation in the experiments above. The addition of spike-in RNA is effective as processing control when working with challenging and low input material, and can be used to normalize sequencing reads or calculate input RNA amounts. In addition, the correlation values between the expected and observed relative quantities of the spikes can be calculated. The high correlation in our experiments indicate excellent recovery of the ERCC spike-in mix during the entire library preparation and sequencing workflow in all samples but the conditioned medium (SupFig5).

As there is an inverse relationship between the number of spike-in RNA reads and the number of endogenous RNA reads, the ratio between the sum of the reads consumed by the endogenous transcripts and the total number of spike-in reads is a relative measure for the RNA concentration of the various samples. When adding the same amount of ERCC RNA to all samples, a higher ratio is indicative of more endogenous RNA. We found the highest RNA extraction concentration in conditioned medium, and the lowest in plasma EVs (SupFig6). Of note, not all starting volumes before EV purifications or other handling were equal. For instance, in our urine experiment we compare RNA extracted from 200 uL whole urine with RNA isolated from EVs that were present in 45 mL whole urine as starting material. Therefore, we corrected the endogenous:ERCC ratios for the original input volumes. This provides us information about the relative amount of RNA present per milliliter biofluid (Fig 3A). While ePRP, conditioned medium and urine have very similar RNA concentrations, ePFP and cPFP contain approximately 17 times less RNA. In addition, EVs from condition medium hold 2763 times less RNA compared to their fluid of origin, plasma EVs 616 times less and urine EVs 7.6 times less. Given that only one biological sample was included in this experiment, further studies warranted to validate these differences in RNA concentration.

**Figure 3.**
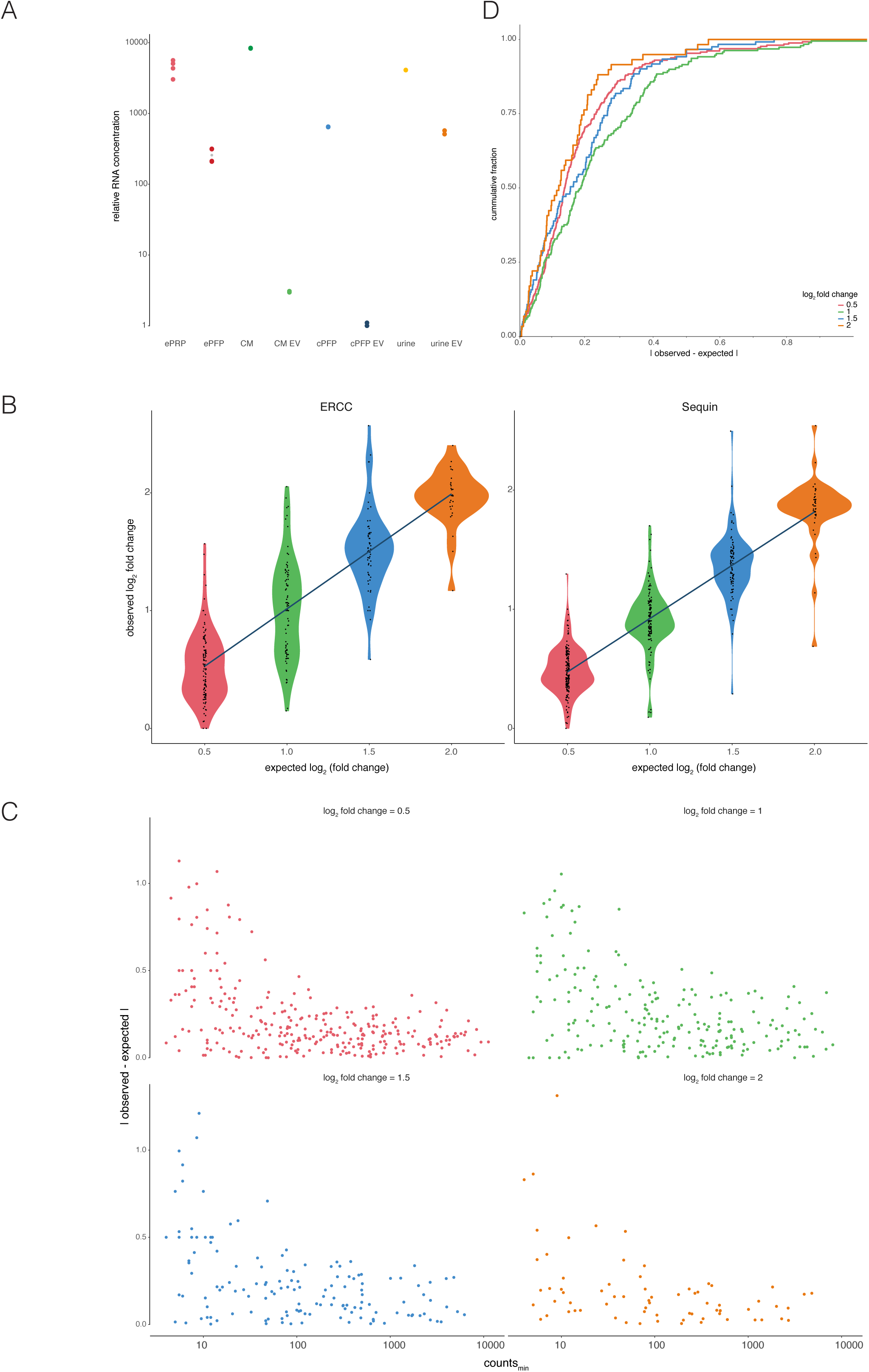
Spike-in RNA based assessment of relative RNA concentration and trueness. A) Relative RNA concentration estimation. B) Relationship between expected and observed log_2_ fold changes shows an overall good correlation. C) The log_2_ fold change differences are higher in spikes with low counts. D) Cumulative distributions of log_2_ fold change differences demonstrate good concordance between expected and observed differences.

In a separate experiment, we added two different spike-in mixes in varying amounts to five identical ePFP samples from a fourth healthy donor. Sequin spikes (n=78) and ERCC spikes (n=92) were diluted in opposite order by a factor 1.41 in the five derivative samples. In this way, a biologically relevant 4-fold dynamic range for both Sequin and ERCC spikes was covered (Fig 1C). The aim of this experiment was to assess the method’s trueness by comparing expected and observed fold changes of the 170 sequenced spike-in RNAs. Of note, both Sequin and ERCC spike mixes consist of multiple RNA molecules present in varying concentrations. Based on pre-experiments, we made sure that we added the spikes in such amounts that the number of reads going to the spikes with the highest concentration (for both the Sequin and ERCC panel) was lower than the number of reads going to the 10^th^ highest abundant endogenous gene. Only by aiming for coverage in the biofluid abundance range, one is able to assess the accuracy of biologically relevant differences. The results indicate how reliably fold changes can be detected using our total RNA seq method. Overall, there is a strong correlation between the expected and observed fold changes, with ERCC spikes (slope=0.975, adjusted R^2^= 0.67) behaving slightly better than Sequin spikes (slope=0.895, adjusted R^2^= 0.78) since the slope is expected to be ‘1’ (Fig 3B). Notably, larger variations arise when assessing smaller fold changes. Indeed, the lower the fold change, the bigger the spread in datapoints in the violin plot. We investigated this observation in more detail and found that deviation from the expected value is larger for spikes with fewer counts (Fig 3C). In order to reliably measure small fold changes, it appears that a minimal number of 10 counts is advisable. Importantly, for about 90% of the spikes the deviation between the observed and expected log_2_ fold change is smaller than 0.5. This is shown in the cumulative distribution plot, where a minimum of 87.3% (for a log_2_ fold change difference of 1) and a maximum of 91.4 % (for a difference of 2) of the spikes show a deviation from the expected value of maximum 0.5 (Fig 3D). This indicates that the worst measurement for about 90% of the spikes is wrong with only a factor 1.41. What is more, almost all spikes can be measured within an error of a factor 2. In conclusion, although very small fold changes and fold changes of lower abundant transcripts are somewhat more difficult to detect, the method is reliable and approximates true fold changes very well.

### 2.3. The total RNA seq method is reproducible

As indicated above, technical replicates of the e PRP and ePFP samples were prepared at the level of RNA isolation. Scatter plots of the read counts clearly show that gene counts are reproducible between independent RNA extractions of the same plasma sample (Fig 4). In addition, we generated cumulative distribution plots that display the fold change of every gene when comparing RNA isolation replicates (Fig 5A). The area left of the curve (ALC) indicates the precision of the method, with lower values demonstrating better replication. Indeed, the more the curves are shifted to the left, the smaller the differences between two replicates and thus the smaller the ALC value. In biological terms, this means that half of the genes can be detected with a fold change smaller than the ALC value. To illustrate, in ePRP of donor 2 half of the genes show a fold change less than 1.32 between both replicates (log_2_ fold change of 0.403, indicated in Fig 5C). Cumulative distribution plots for the experiment with conditioned medium, citrate plasma, urine and their respective EVs (Fig 5B). show slightly lower ALC values, indicating that reproducibility is better when replication is introduced at the level of library preparation (Fig 5C).

**Figure 4.**
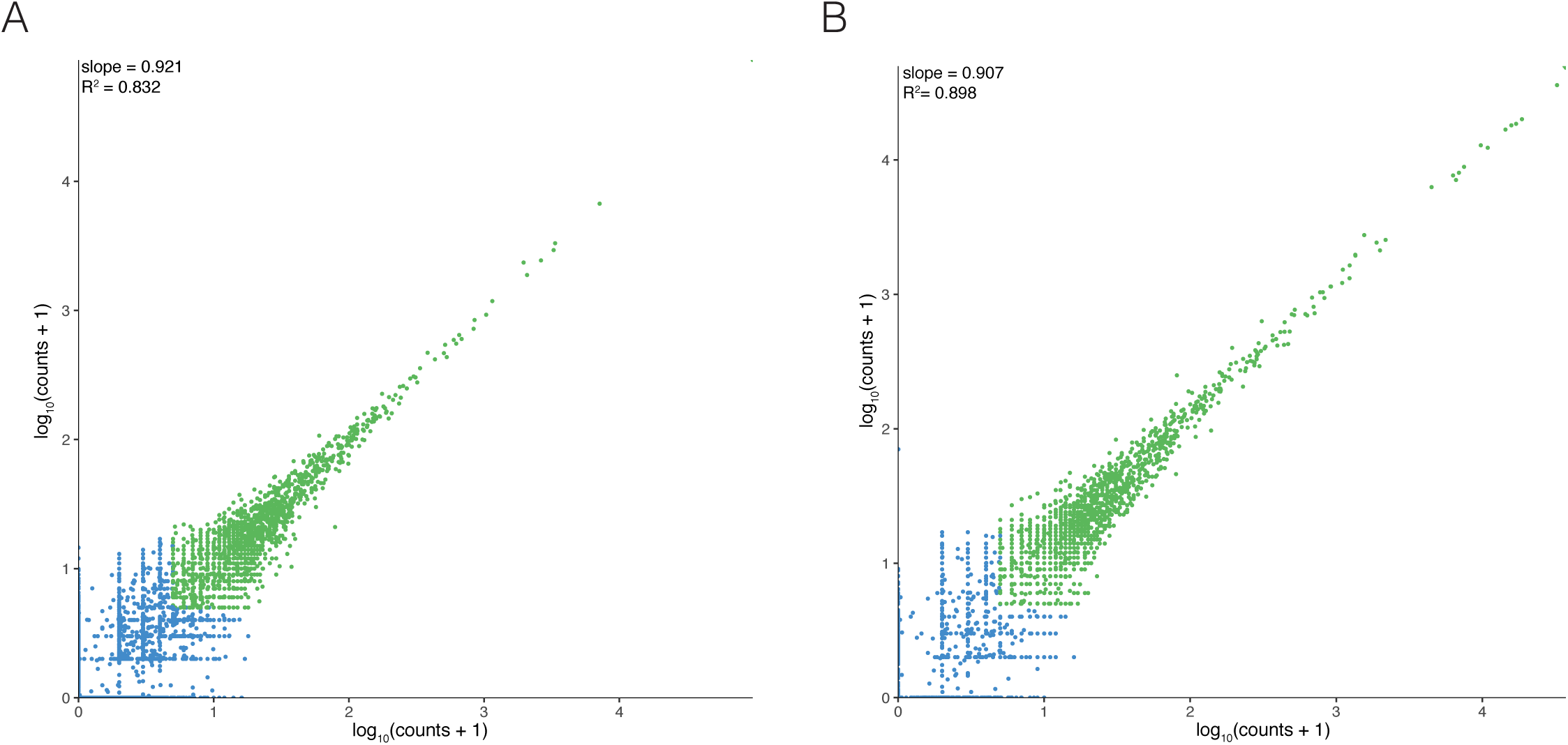
RNA isolation replicates of ePRP and ePFP show high repeatability. A) ePRP and B) ePFP replicate correlation with filtered (counts < 4, red) and retained genes (counts >= 4, green) resulted in high Pearson correlation of 0.912 and 0.948, respectively.

**Figure 5.**
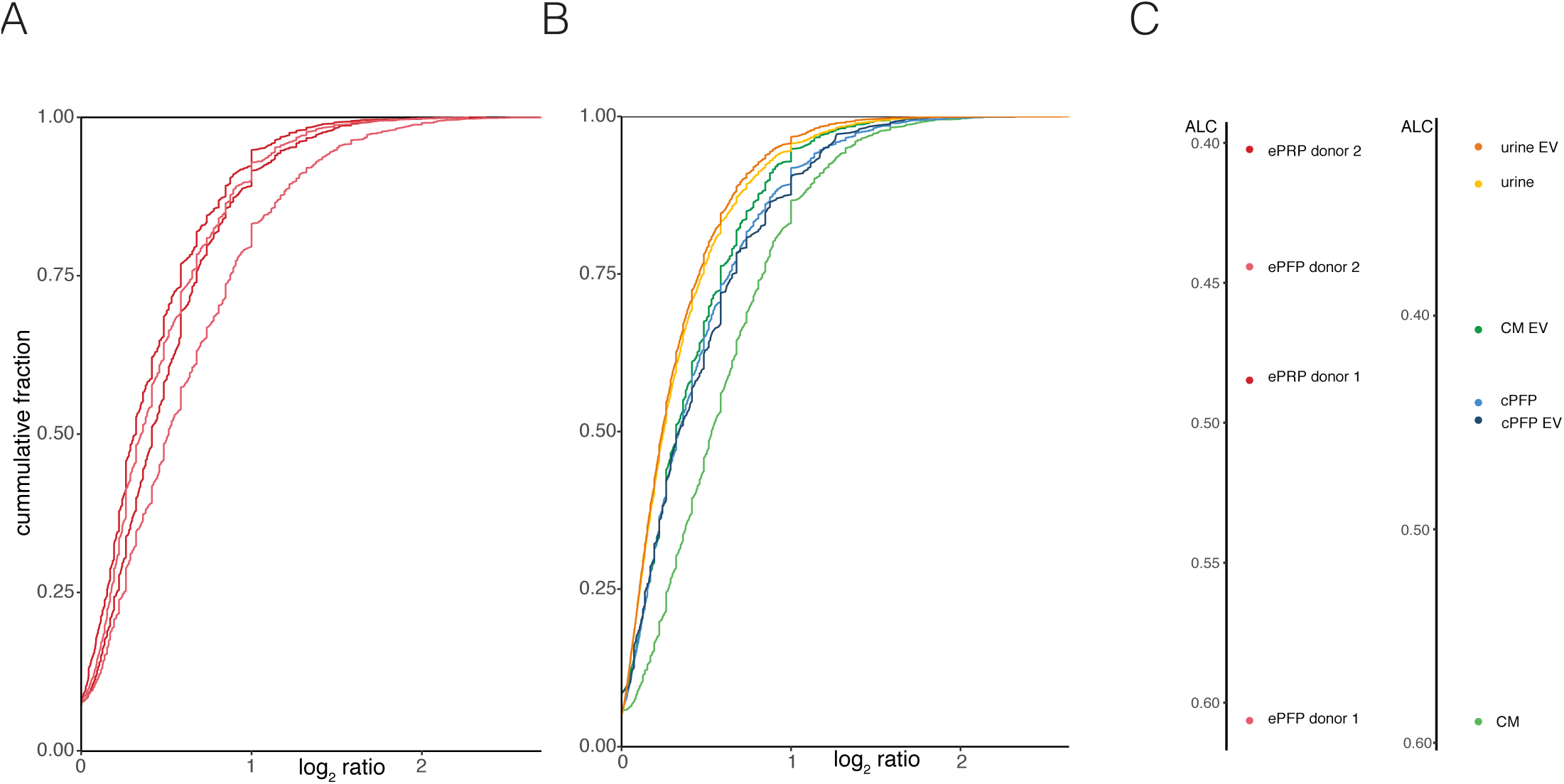
Cumulative distributions of the log_2_ ratio for all replicate pairs with their respective values of the area left of the curve. A) ePRP and ePFP RNA isolation replicates of two donors. B) Library preparation replicates of CM, CM-EV, cPFP, cPFP-EV, urine and urine-EV.

### 2.4. Transcriptomes are widely different among tested biofluids

To assess the inherent variation of the various transcriptomes, we clustered all plasma, urine, conditioned medium and EV samples in a t-SNE plots (SupFig7). This plot confirms good reproducibility among technical replicates. Notably, EVs isolated from healthy donor plasma and cancer cell conditioned medium seem to be quite similar. In contrast, urinary EVs do not cluster with these EVs, but show more similarity to whole urine. Next, when assessing the number of reproducibly detected genes (mRNA, lncRNA, miscRNA pseudogenes and others), ePFP samples contain more genes compared to ePRP (Fig 6A). This is probably due to lower amounts of (very abundant) mitochondrial RNA in ePFP, hence freeing up sequencing power to detect more genes. In addition, the 20 most abundant genes consume approximately 75% of the reads in ePRP, automatically leading to less diversity in the remaining gene fraction (Fig 6B). The highest abundant genes in PRP are MTRNR2 (or paralogues), MTND1 and MTND2, which are all transcribed from mitochondrial DNA, as are many other genes in the top-20 (SupFig8). Urine and urinary EVs contain more than 10,000 genes in our experimental setup, the highest number of all evaluated biofluids (Fig 6C). The lowest number of genes was observed in healthy donor citrate plasma derived EVs, in which only 904 genes could be detected using our total RNA seq method. Interesting to note is that plasma EVs had the worst mapping qualities of all samples (see Fig 2A above). An important remark is that one should be cautious when interpreting the results above. Indeed, simply comparing gene numbers among different biofluids is difficult because of varying input volumes used for RNA purification. As already exemplified above, in the urine experiment we compare RNA extracted from 200 uL whole urine with RNA isolated from EVs that were present in 45 mL whole urine as starting material. To get further insights in the technical performance of the total RNA seq method, we also assessed the distribution of the counts (SupFig9) and the gene body coverage (SupFig10). In fragmented RNA, the coverage at the 5’ and 3’ end of the gene body is typically lower compared to the middle part.

**Figure 6.**
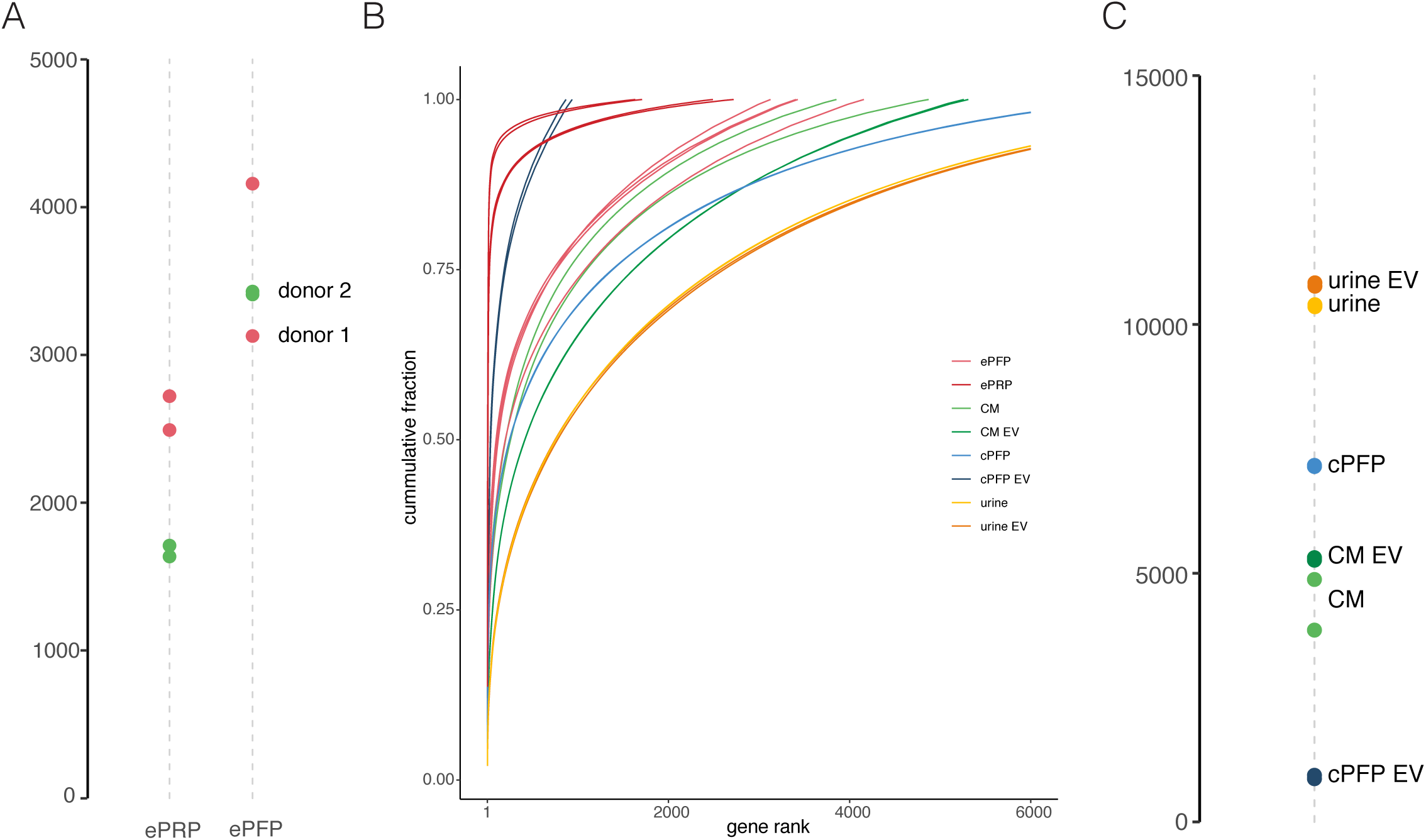
The number of genes differs among sample types. A) Number of genes (counts >=4) detected in ePRP and ePFP. B) Read consumption of the genes ranked by abundance. B) Number of genes (counts >=4) detected in CM, CM-EV, cPFP, cPFP-EV, urine and urine-EV.

We further investigated five different gene biotypes in all samples, according to their annotation in Ensembl (protein coding genes, lncRNA genes, miscellaneous RNA genes, pseudogenes and other genes). The percentage of counts assigned to these five gene types differs among the biofluids. ePRP for instance contains high number of pseudogene reads, resulting from mitochondrial genes as illustrated above, whereas ePFP mainly consists of reads mapping to protein coding genes (Fig 7A). The differences in the other samples are less explicit. Looking into the top-20 genes with the highest counts reveals the genes consuming most of the reads in each sample (SupFig8). We also calculated the absolute numbers per gene biotype, but again we should keep in mind the difficulty in side-by-side comparisons because of differing input volumes (Fig 7B-C). What we can conclude is that the method is able to pick up many different classes of RNA molecules.

**Figure 7.**
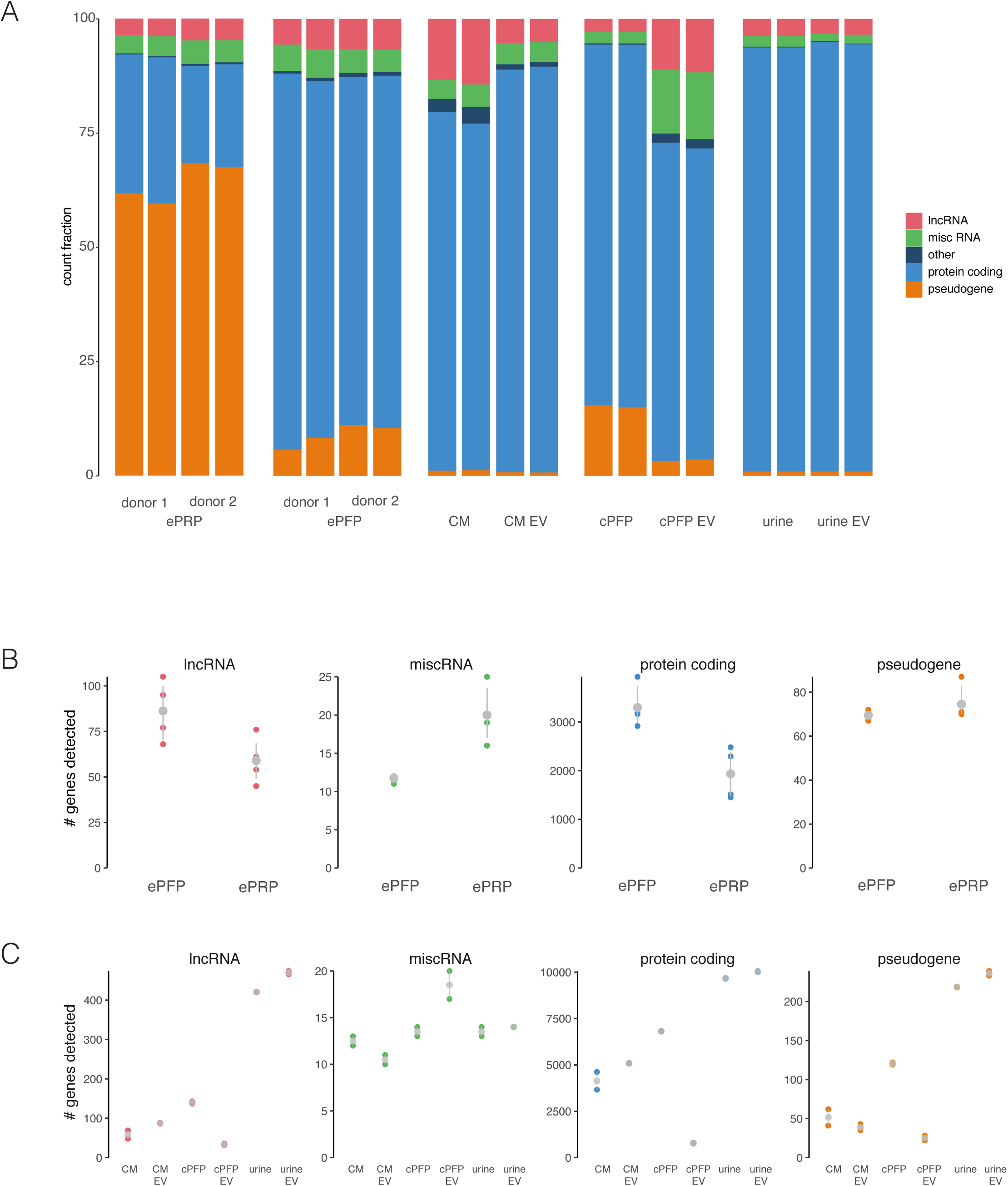
Detected gene-biotypes differ among sample types. A) Percentage of exonic reads attributed to the different biotypes per sample quantified with Kallisto. B-C) Detected number of genes per biotype for all samples.

Next to Ensembl, we also assessed the reads mapping to LNCipedia^11^, the most comprehensive database of human long non-coding RNAs (Fig 8A). In analogy with the results above, the largest number of lncRNAs was found in urine and urinary EVs. Indeed, approximately 3000 lncRNA genes can be distinguished in EVs isolated from urine. cPFP contains around 1500 lncRNAs, while we could detect almost no lncRNAs in EVs isolated from this plasma. As expected, ePFP contains more lncRNAs than ePRP. In addition, also the presence of circular RNAs was assessed. Their overall number is low, but especially cPFP and urinary EVs show substantially more circular RNAs (Fig 8B). CircRNAs are presumed to be more stable and less degraded compared to linear forms. Therefore, they are ideal candidates for cancer biomarker discovery studies.

**Figure 8.**
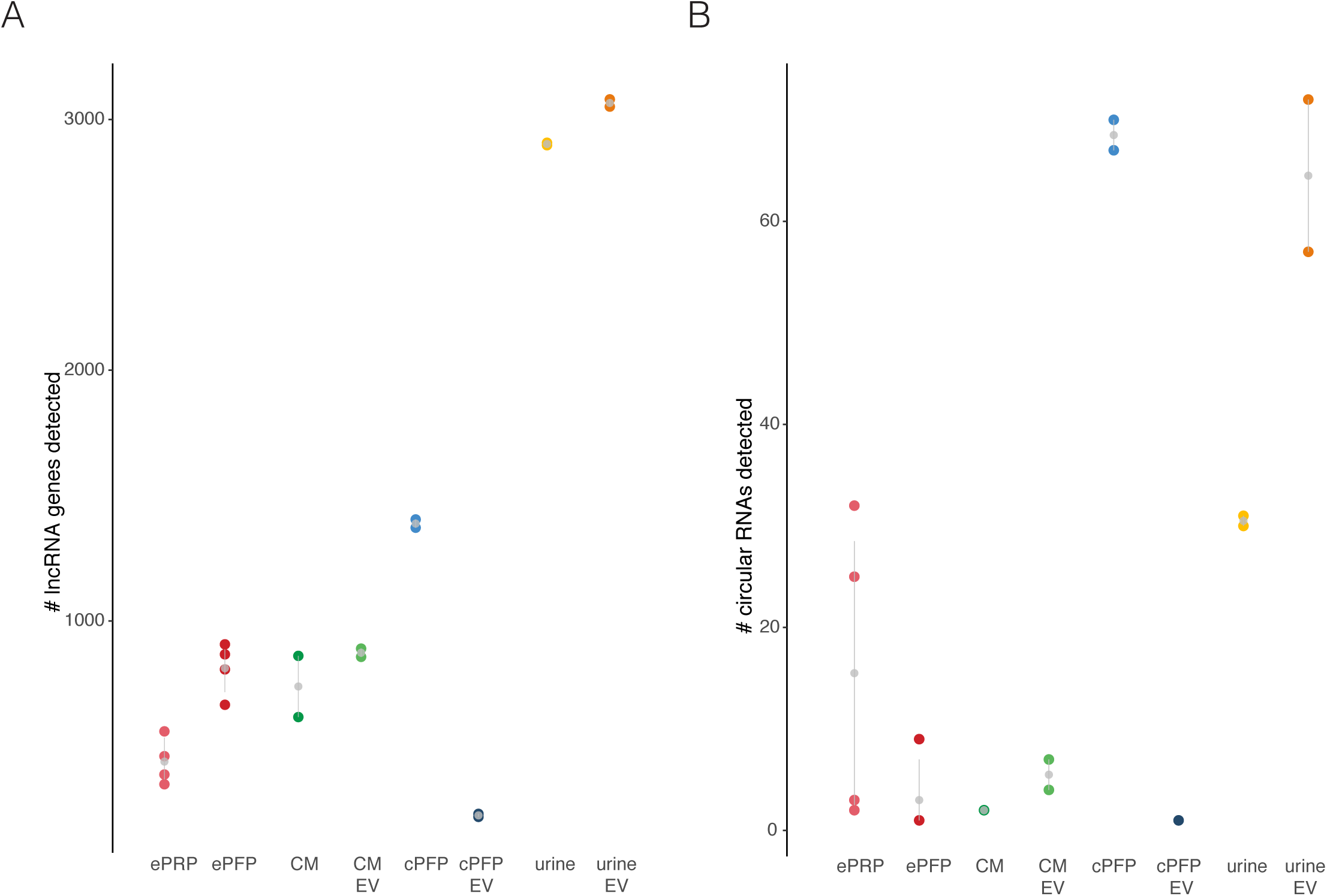
Non-coding RNAs, both linear and circular, are detected in total RNA sequencing libraries. A) Number of lncRNAs quantified based on LNCipedia. B) Number of circular RNAs detected with CircExplorer2.

### 2.5. Evaluating biological differences in RNA content among biofluids

In order to illustrate which biological insights total RNA seq results can yield, we compared gene abundance in ePRP and ePFP samples (SupFig11A). An Euler diagram indicates the number of genes that are unique to each plasma fraction, and the number of overlapping genes (SupFig11B). Studies like this (but with many more samples in each biofluid group) could lead to new insights into selective RNA cargo filling of extracellular vesicles. Here, we compared RNA abundance profiles between EVs and their biofluids of origin. Euler diagrams represent the number of overlapping and unique genes per pair of samples (Fig 9A-C). Conditioned medium, for instance, shares 4891 genes (Jaccard index of 0.652) with the EVs it contains. Further, 1853 genes are only present in EVs while 755 genes are unique to conditioned medium only. The results in plasma are markedly different: plasma EVs contain 1598 genes, 70 of which are unique to EVs. RNA isolated from whole citrate plasma on the other hand contains 7211 genes, nearly five times more, despite 30-fold lower input volume. Urine and urinary EVs finally have more than 10,000 genes in common and contain 521 and 900 unique genes respectively. In addition, using scatter plots we represent the similarity between abundances in EVs and their fluid of origin in another way (Fig 9D-E). Supporting the results above, urine and urinary EVs have a great concordance in abundance of genes while citrate plasma and plasma EVs differ most from each other. Note that most of the EV-unique genes (indicated with dark blue dots) are low abundant. This could be due to chance (sampling effect) and sequencing deeper or using more input material may reduce this set of unique genes. In the same plot, we also indicated the count level of all genes uniquely present in one of both samples with colored lines. Notably, genes present in EVs but absent from their biofluid of origin typically consume a lower number of counts. Digging deeper into biological analyses using bigger cohorts, from gene set enrichment to pathway analysis, may reveal novel insights.

**Figure 9.**
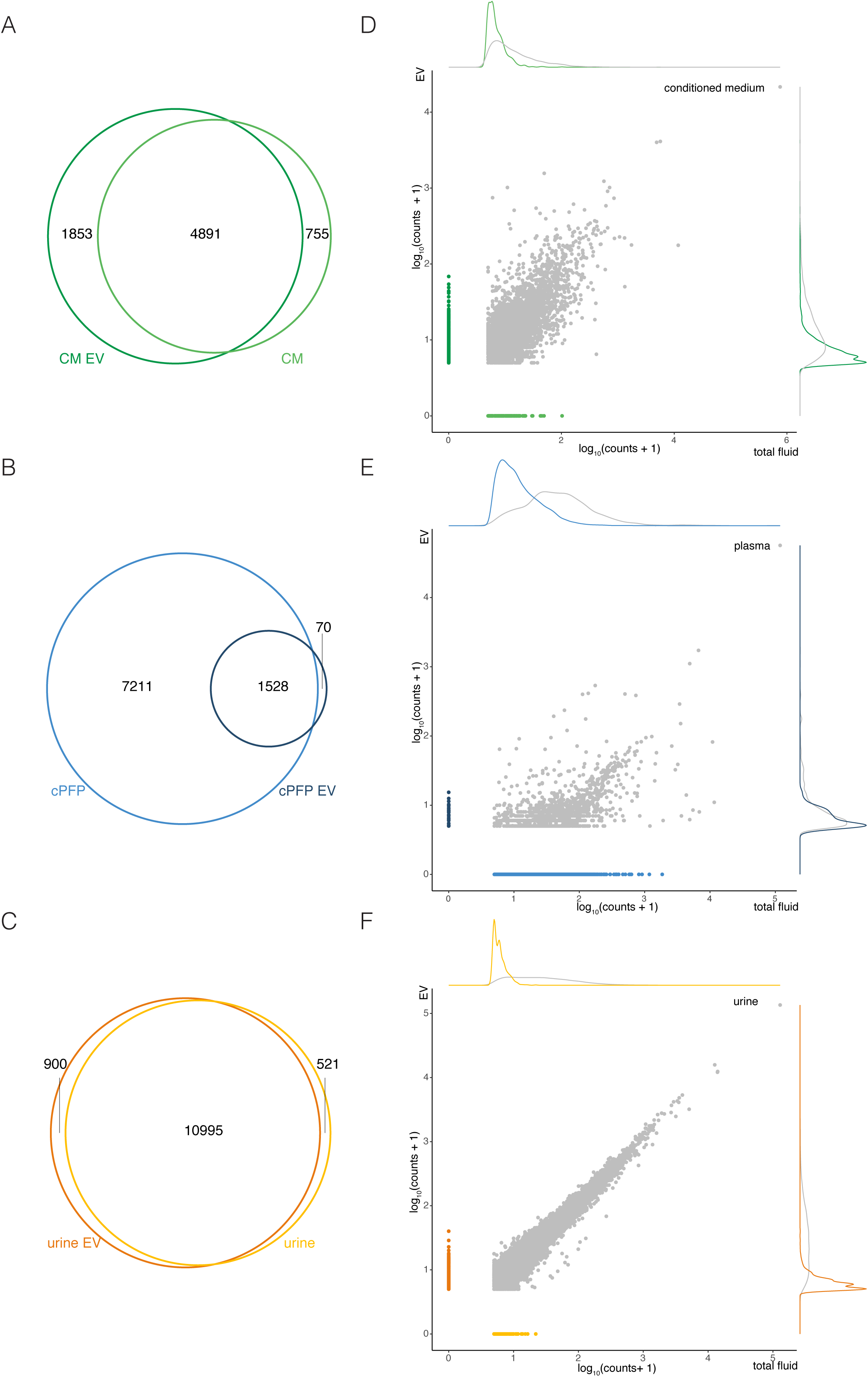
Gene detection overlap and correlation between EVs and their biofluid of origin differ among the sample biotypes. Euler diagrams of A) CM and CM-EV, B) cPFP and cPFP-EV, and C) urine and urine-EV. Correlation of overlapping (gray) and specific genes (colored) between EVs and their origin for D) CM and CM-EV, E) cPFP and cPFP-EV, and F) urine and urine-EV.

## 3. Discussion

Extracellular RNA content analysis of human biofluids and extracellular vesicles may provide insights into their biogenesis and reveal biomarkers for health and disease. There are currently four types of sequencing-based total RNA profiling of such challenging clinical samples: 1) the recent modified small RNA sequencing methods^8,9^, 2) the SOLiD total RNA sequening method^12^, 3) the Ion Proton method^13^ and 4) TGIRT-sequencing using thermostable group II intron reverse transcriptases^5^. The SMARTer method assessed in our study adds a fifth promising method to the sequencing armory. In addition, the SMARTer method avoids limitations linked to other methods such as short fragment length, low amount of quantified genes or ribosomal RNA contamination.

While not marketed for this application, extensive technical performance assessment demonstrated that the SMARTer Stranded Total RNA-Seq method to be an accurate, precise and sensitive method to quantify total RNA in human biofluids. Notable differences among plasma, urine, conditioned medium and their EVs could be related to the biology of each fluid and should be taken into account when setting up biomarker studies. Possible improvements to profile platelet-rich plasma from EDTA tubes could be made by designing probes that remove mitochondrial ribosomal RNA, shown to be highly abundant (and unwanted) in this type of plasma. In this way, read diversity should increase and more genes at lower abundance will be identified. Quite striking was the observation that EVs from platelet-free citrate plasma contain substantially fewer genes. Whether the workflow can be optimized for plasma EVs definitely is a subject for further research. Besides, treatment of EVs with RNases to remove any non-encapsulated RNA may also prove useful^14^.

It has been shown that pre-analytical variables may have an effect on the resulting RNA profiles^15^. In our study, we also observed differences between ePFP and cPFP, which are identical biofluids collected in different blood tubes and prepared with a slightly different centrifugation protocol. In general, differences in pre-analytical variables such as blood collection tubes, processing time, centrifugation speeds, RNA isolation kit, and freeze-thaw cycles could well be responsible for great variation in RNA sequencing results. Systematic evaluation of the impact of pre-analytical variables would definitely be of huge added value to progress the fields of extracellular RNA research and liquid biopsies.

In our study we included synthetic spike-in RNA mixes to control for variation during RNA isolation and/or library preparation. Of note, we did not include spikes during RNA isolation of EVs and their biofluids or origin because we did not include replicates at the RNA level. Ideally however, both Sequin spikes^16^ during RNA extraction and ERCC spikes before library preparation are added in all RNA sequencing experiments to control for different types of technical variation. As data interpretation is often complex in experiments involving different biofluids and input volumes, spike-in RNA could help with normalization, clarification and assimilation of raw data.

Finally, nuclear acids present in all sorts of biofluids and their EVs are promising biomarkers for diagnosis, prognosis, therapy response and monitoring of disease. The advantage of the SMARTer Stranded Total RNA-Seq method is its potential to process low amounts of input material. Indeed, collecting samples is often the bottleneck of fundamental, (pre)clinical and translational research projects and being able to disseminate large amounts of information from only 200 μL (or less) can substantially impact research progress.

## 4. Materials and Methods

### 4.1 Sample collection

#### 4.1.1. ePRP and ePFP collection

For the first experiment, venous blood was drawn from an elbow vein of two healthy donors in 3 EDTA tubes (BD Vacutainer Hemogard Closure Plastic K2-Edta Tube, 10 ml, #367525) using the BD Vacutainer Push blood collection set (21G needle). Collection of blood samples was according to the Ethical Committee of Ghent University Hospital approval EC/2017/1207 and written informed consent of the donors was obtained. The tubes were inverted 5 times and centrifuged within 15 minutes after blood draw (400 g, 20 minutes, room temperature, without brake). Per donor, the upper plasma fractions were pipetted (leaving approximately 0.5 cm plasma above the buffy coat) and pooled in a 15 ml tube. After gently inverting, five aliquots of 220 µl platelet-rich plasma (ePRP) were snap-frozen in 1.5 ml LoBind tubes (Eppendorf Protein LoBind microcentrifuge tubes Z666548 - DNA/RNA) in liquid nitrogen and stored at −80 °C. The remaining plasma was centrifuged (800 g, 10 minutes, room temperature, without brake) and transferred to a new 15 ml tube, leaving approximately 0.5 cm plasma above the separation. This plasma was centrifuged a 3^rd^ time (2500 g, 15 minutes, room temperature, without brake), and transferred to a 15 ml tube, leaving approximately 0.5 cm above the separation. The resulting platelet-free plasma (ePFP) was gently inverted, snap-frozen in five aliquots of 220 µl and stored at −80 °C. The entire plasma preparation protocol was finished in less than two hours. 200 µl ePRP and ePFP was used for each RNA isolation. For the spike-in RNA titration experiment, the protocol was identical except for the fact that 4 EDTA tubes of 10 ml were drawn and that the second centrifugation step was different (1500 g, 15 minutes, room temperature, without brake).

#### 4.1.2 cPFP collection and EV isolation

Venous blood was collected using a 21G needle in 3.2% (w/v) sodium citrate tubes (MLS, Menen, Belgium) from an elbow vein of a healthy donor. Collection of blood samples was according to the Ethical Committee of Ghent University Hospital approval EC/2014/0655 and in accordance to relevant guidelines. The participant had given written informed consent. Absence of hemolysis was confirmed by the lack of a spectrophotometric absorbance peak of free hemoglobin at 414 nm using a BioDrop DUO spectrophotometer (BioDrop Ltd, Cambridge, United Kingdom). The blood tubes were inverted 5 times and plasma was prepared by centrifugation (2500 g with brake, 15 minutes, room temperature). The upper plasma fraction was collected (leaving approximately 0.5 cm plasma above the buffy coat layer) and transferred to a new 15 ml tube. Platelet-depleted plasma was prepared by centrifugation (2500 g with brake, 15 minutes, room temperature). Platelet-depleted plasma was collected (leaving approximately 0.5 cm plasma above the bottom of the tube), aliquoted per 1.5 ml in 2 ml cryo-vials and stored at −80 °C. To ensure the depletion of platelets in plasma we used the XP-300 Hematology Analyzer (Sysmex, Hoeilaart, Belgium). The blood sample was processed within 120 min after blood collection. 200 µl plasma was used for RNA isolation.

A combination of size exclusion chromatography (SEC) and OptiPrep density gradient (DG) centrifugation was used to isolate EV from plasma. Sepharose CL-2B (GE Healthcare, Uppsala, Sweden, #17014001) was washed 3 times with PBS (Merck Millipore, Billerica, Massachusetts, USA) containing 0.32 % (w/v) trisodiumcitrate dihydrate (ChemCruz, Dallas, Texas, USA)^17^. For preparation of the SEC column, nylon filter with 20 µm pore size (Merck Millipore, Billerica, Massachusetts, USA) was placed on bottom of a 10 ml syringe (Romed, Wilnis, The Netherlands), followed by stacking of 10 ml Sepharose CL-2B. On top of three SEC columns, 6 ml plasma was loaded (2 ml per column) and fractions of 1 ml eluate were collected. SEC fractions 4, 5 and 6 were pooled and concentrated to 1 ml using 10 kDa centrifugal filter (Amicon Ultra-2ml, Merck Millipore, Billerica, Massachusetts, USA). The resulting 1 ml sample was loaded on top of a DG, as previously described^18^. This discontinuous iodixanol gradient was prepared by layering 4 ml of 40 %, 4 ml of 20 %, 4 ml of 10 % and 3.5 ml of 5 % iodixanol in a 17 ml Thinwall Polypropylene Tube (Beckman Coulter, Fullerton, California, USA). The DG was centrifuged 18 h at 100,000 g and 4 °C using SW 32.1 Ti rotor (Beckman Coulter, Fullerton, California, USA). Density fractions of 1 ml were collected and fractions 9-10 pooled. An additional SEC was performed on the pooled density fraction to remove iodixanol^19^. SEC fractions 4-7 were pooled and concentrated to 100 µl and stored at −80 °C until further use. Samples were further diluted to 200 µl in PBS prior to RNA isolation.

#### 4.1.3 Urine collection and EV isolation

One whole urine sample was collected from a prostate cancer patient prior to local treatment. Sample collection was according to the Ethical Committee of Ghent University Hospital approval EC/2015/0260 and in accordance to relevant guidelines. The participant had given written informed consent. The urine sample was collected immediately following digital rectal examination (DRE). DRE was performed as 3 finger strokes per prostate lobe. The urine sample was centrifuged for 10 minutes at 1000 g and 4 °C in accordance with the Eurokup/HKUPP Guidelines. Cell-free urine supernatants were collected (leaving approximately 0.5 cm urine above the cell pellet) and stored at −80 °C in 1.7 ml SafeSeal Microcentrifuge Tubes (Sorenson Bioscience) until further use. 200 µl urine was used for RNA isolation.

The cell-free urine sample (45 ml) was thawed at room temperature and vortexed extensively before being concentrated to 800 µl using a 10 kDa centrifugal filter device (Centricon Plus-70, Merck Millipore, Massachusetts, USA). The concentrated urine sample was resuspended in 3.2 ml of a 50% iodixanol solution and layered on the bottom of a 17 ml Thinwall Polypropylene Tube (Beckman Coulter, Fullerton, California, USA). A discontinuous DG was prepared by additional layering of 4 ml of 20%, 4 ml of 10% and 3.5 ml of 5% iodixanol, and 1 ml PBS on top of the urine suspension. The DG was centrifuged 18 h at 100,000 g and 4 °C using SW 32.1 Ti rotor (Beckman Coulter, Fullerton, California, USA). Density fractions of 1 ml were collected and fractions 9-10 pooled. An additional SEC was performed on the pooled density fraction to remove iodixanol. SEC fractions 4-7 were pooled and concentrated to 100 µl and stored at −80 °C until further use. Samples were further diluted in PBS to 200 µl for RNA isolation.

#### 4.1.4 MCF-7 GFP-Rab27b conditioned medium and EV isolation

The MCF-7 cell line (ATCC, Manassas, VA, USA) was stably transfected with peGFP-C1 vector (Clontech, Mountain View, California, USA) containing the GFP-Rab27b fusion protein, as previously described (MCF-7 GFP-Rab27b)^20^. MCF-7 GFP-Rab27b cells were cultured in Dulbecco’s Modified Eagle Medium supplemented (DMEM) with 10 % fetal bovine serum, 100 U/ml penicillin, 100 µg/ml streptomycin and 1 mg/ml G418. Presence of mycoplasma was routinely tested using MycoAlert Mycoplasma Detection Kit (Lonza, Verviers, Belgium). To prepare conditioned medium (CM), 4 x 10^8^ MCF-7 GFP-Rab27b cells (20 × 175 cm^2^ flasks, 300 ml) were washed once with DMEM, followed by two washing steps with DMEM supplemented with 0.5 % EV-depleted fetal bovine serum (EDS). EDS was obtained after 18 h ultracentrifugation at 100,000 g and 4 °C (SW55 Ti rotor, Beckman Coulter, Fullerton, California, USA), followed by 0.22 µm filtration. Flasks were incubated at 37 °C and 10 % CO_2_ with DMEM containing 0.5% EDS. After 24 h, CM was collected and centrifuged for 10 min at 200 g and 4 °C. Cell counting was performed with trypan blue staining to assess cell viability (Cell Counter, Life Technologies, Carlsbad, California, USA). The supernatant was passed through a 0.45 µm cellulose acetate filter (Corning, New York, USA) and CM was concentrated to 1 ml at 4 °C using a 10 kDa Centricon Plus-70 centrifugal unit (Merck Millipore, Billerica, Massachusetts, USA). 200 µl was used for RNA isolation. After filtering through a 0.22 µm filter (Whatman, Dassel, Germany), 1 ml concentrated conditioned medium (CCM) was used for DG ultracentrifugation. Fractions of 1 ml were collected and fractions 9-10 pooled. Pooled fractions were diluted to 15 ml with phosphate-buffered saline (PBS), followed by 3 h ultracentrifugation at 100,000 g and 4 °C using SW 32.1 Ti rotor (Beckman Coulter, Fullerton, California, USA). Resulting pellets were resuspended in 100 µl PBS and stored at −80 °C until further use. Samples were further diluted in PBS to 200 µl for RNA isolation.

### 4.2 Extracellular vesicle quality control

We have submitted all relevant data of our experiments to the EV-TRACK knowledgebase^21^ (EV-TRACK ID: EV190039).

#### 4.2.1 Antibodies

The following antibodies were used for immunostaining: anti-Alix (1:1000, 2171S, Cell Signaling Technology, Beverly, Massachusetts, USA), anti-TSG101 (1:1000, sc-7964, Santa Cruz Biotechnology, Dallas, Texas, USA), anti-CD9 (1:1000, D3H4P, Cell Signaling Technology, Beverly, Massachusetts, USA), anti-THP (1:800, sc-20631, Santa Cruz Biotechnology, Dallas, Texas, USA), anti-Flot-1 (1:1000, 610820, BD Biosciences, Franklin Lakes, New Jersey, USA), anti-Ago2 (1:1000, ab32381, Abcam, Cambridge, UK), anti-ApoA-1 (1:100, B10, Santa Cruz Biotechnology, Dallas, Texas, USA), sheep anti-mouse horseradish peroxidase-linked antibody (1:3000, NA931V, GE Healthcare Life Sciences, Uppsala, Sweden), donkey anti-rabbit horseradish peroxidase-linked antibody (1:4000, NA934V, GE Healthcare Life Sciences, Uppsala, Sweden).

#### 4.2.2 Protein analysis

EV protein concentrations were measured using the fluorometric Qubit Protein Assay (ThermoFisher, Waltham, Massachusetts, USA). Sample preparation was done by 1:1 dilution with SDS 0.4%. Protein measurements were performed using the Qubit Fluorometer 3.0 (ThermoFisher, Waltham, Massachusetts, USA) according to the manufacturer’s instructions.

ODG fractions were dissolved in reducing sample buffer (0.5 M Tris-HCl (pH 6.8), 40% glycerol, 9.2% SDS, 3% 2-mercaptoethanol, 0.005% bromophenol blue) and boiled at 95 °C for 5 min. Proteins were separated by SDS-PAGE (SDS-polyacrylamide gel electrophoresis), transferred to nitrocellulose membranes (Bio-Rad, Hercules, California, USA), blocked in 5% non-fat milk in PBS with 0.5% Tween-20 and immunostained. Chemiluminescence substrate (WesternBright Sirius, Advansta, Menlo Park, California, USA) was added and imaging was performed using the Proxima 2850 Imager (IsoGen Life Sciences, De Meern, The Netherlands).

#### 4.2.3 Nanoparticle tracking analysis

EV samples were analyzed by Nanoparticle tracking analysis (NTA) using a NanoSight LM10 microscope (Malvern Instruments Ltd, Amesbury, UK) equipped with a 405 nm laser. For each sample, three 60 second videos were recorded at camera level 13. Temperature was monitored during recording. Recorded videos were analyzed at detection threshold 3 with NTA Software version 3.2 to determine the concentration and size distribution of measured particles with corresponding standard error. For optimal measurements, samples were diluted with PBS until particle concentration was within the optimal concentration range for particle analysis (3×10^8^-1×10^9^).

#### 4.2.4 Transmission electron microscopy

EV samples were qualitatively and quantitatively analyzed with transmission electron microscopy (TEM). Samples were deposited on Formvar carbon-coated, glow discharged grids, stained with uranylacetate and embedded in methylcellulose/uranylacetate. These grids were examined using a Tecnai Spirit transmission electron microscope (FEI, Eindhoven, The Netherlands) and images were captured with a Quemasa charge-coupled device camera (Olympus Soft Imaging Solutions, Munster, Germany).

### 4.3 RNA isolation, spike-in RNA addition and DNase treatment

RNA isolation was performed using the miRNeasy Serum/Plasma Kit (Qiagen). In experiment 1, ePRP and ePFP RNA was isolated from 200 µl of platelet-rich and platelet-free plasma from two healthy donors. Two RNA replicates were included. 2 µl of Sequin RNA spikes^16^ were added to the lysate at a dilution of 1/3000 for PFP and 1/250 for PRP, to control for variation in RNA isolation. After isolation, 2µl of ERCC RNA spikes (ThermoFisher) were added to the eluate at a dilution of 1/25 000 for PFP and 1/5000 for PRP. This allows to estimate the relative concentration of the eluate. For the ePFP RNA of the healthy donor, used for the spike-in RNA titration experiment (see 4.4), we used 6 aliquots of 200 µl plasma and pooled the RNA after isolation. We did not add Sequin spikes during RNA isolation. ERCC spikes were added following a titration series, as described in the next paragraph. Finally, RNA from EVs and their respective biofluids was isolated with the same kit, using 200 µl sample input (see also 4.1). No duplicates were included at the level of RNA isolation, no Sequin spikes were added, and the standard spin columns were replaced by Ultra-Clean Production (UCP) columns (Qiagen). ERCC spikes were added to the RNA isolation eluate at a dilution of 1/30 000 for plasma and urine and 1/50 for conditioned medium.

### 4.4 Spike-in RNA titration for assessment of trueness

Pooled ePFP RNA (prepared without Sequin spike-in RNA addition) was distributed in five separate tubes, each containing 12 µl RNA. Then, we added 1 µl DNase, 1.6 µl reaction buffer, 2 µl Sequin spikes and 2 µl ERCC spikes to each tube. Both spike-in RNA types were added in a 5-point 1.414-fold dilution series, in opposing order. For Sequin: 1/15,000, 1/21,277, 1/30,000, 1/42,433 and 1/60,000. For ERCC: 1/100,000, 1/70,721, 1/50,000, 1/35,461 and 1/25,000.

### 4.5 Total RNA library preparation and sequencing

On the total amount of 12 µl eluate, gDNA heat-and-run removal was performed by adding 1 µl of HL-dsDNase (ArcticZymes 70800-202, 2 U/µl) and 1 µl reaction buffer (ArcticZymes 66001). Of the resulting volume, 4 µl was used as input for the total RNA library preparation protocol. Sequencing libraries were generated using SMARTer Stranded Total RNA-Seq Kit v2 - Pico Input Mammalian (Takara, 634413). Compared to the manufacturer’s protocol, the fragmentation step was set to 4 min at 94 °C, hereafter the option to start from highly degraded RNA was followed. Library quality control was performed with the Fragment Analyzer high sense small fragment kit (Agilent Technologies, sizing range 50 bp-1000 bp). Based on Qubit concentration measurements or KAPA qPCR, samples were pooled and loaded on the NextSeq 500 (Illumina) with a loading concentration of 1.1 or 1.2 pM. Note that the 1.2 pM resulted in lower quality reads as the run was slightly overloaded. Paired end sequencing was performed (2×75 bp) with median depth of 15.3 million reads per sample. The fastq data is deposited in GEO (GSE131689).

### 4.6 Sequencing data quality control

The reads with a low quality score (Q30) were discarded, hereafter read duplicates were removed with Clumpify (BBMap v.37.93, standard settings). The libraries were trimmed using cutadapt (v.1.16)^22^ to remove 3 nucleotides of the 5’ end of read 2. To enable a fair comparison, we started data-analysis from an equal number of reads by subsampling to 1 million trimmed and deduplicated reads. To assess the quality of the data, the reads were mapped using STAR (v.2.5.3)^23^ on the hg38 genome including the full ribosomal DNA (45S, 5.8S and 5S) and mitochondrial DNA sequences. The parameters of STAR were according to the ENCODE project. Using SAMtools (v1.6)^24^,reads mapping to the different nuclear chromosomes, mitochondrial DNA and rRNA were extracted and annotated as exonic, intronic or intergenic. The SMARTer total RNA sequencing data is stranded and processed accordingly, so strandedness was considered for each analysis step. Gene body coverage was calculated using the full Ensembl (v91)^25^ transcriptome. The coverage per percentile was calculated.

### 4.7 Quantification of Ensembl and LNCipedia genes, differential abundance analysis and gene set enrichment analysis

Genes were quantified by Kallisto (v.0.43.1)^26^ using both Ensembl (v.91)^25^ extended with the ERCC spike and Sequin spike sequences and LNCipedia (v.5.0)^11^. The strandedness of the total RNA-seq reads was considered by running the –rf-stranded mode. Further processing was done with R (v.3.5.1) making use of tidyverse (v.1.2.1). A cut-off for filtering noisy genes was set based on an analysis of single positive and double positive genes. For a cut-off of 4 counts, at least 95% of the single positive values are filtered out. To measure the biological signal, we first performed differential expression analysis between the treatment groups using DESeq2 (v.1.20.0)^27^. To identify enriched gene sets a fsgea (v.1.6.0) analysis was performed, calculating enrichment for the gene sets retrieved from MSigDB (v.6.2).

### 4.8 Circular RNA detection

CircRNAs were annotated by using the combination of STAR (v.2.6.0)^23^ and CIRCexplorer2 (v2.3.3)^28^. The settings of STAR (used according to Vo et al.) are slightly different compared to linear mapping^4^. Human genome hg38 was used for circRNA analysis. CircRNAs were annotated with host gene names from RefSeq.

## Supplementary Materials

### Supplemental Figures

*Supplemental Figure 1 Read duplication levels are markedly different among different biomaterials*.

*Supplemental Figure 2a Characterization of EV from urine and plasma samples*. Proteins are analyzed by western blot using specific EV markers (ALIX, tsg101, CD9 and flotillin-1) and non-EV markers (THP and ApoA-1). EV samples (density gradient fractions 9-10) are enriched in EV proteins and depleted for contaminants. EVs were qualitatively and quantitatively analyzed by electron microscopy and nanoparticle tracking analysis.

*Supplemental Figure 2b Characterization of EV from MCF-7 GFP-Rab27b cells*. Proteins are analyzed by performing western blot using specific EV markers (ALIX, tsg101 and CD9) and non-EV markers (Ago2). EV samples (density gradient fractions 9-10) are enriched in EV proteins and depleted for contaminants. EVs were qualitatively and quantitatively analyzed by electron microscopy and nanoparticle tracking analysis.

*Supplemental Figure 3. Percentage of reads originating from the sense strand to demonstrate good strandedness of the kit*.

*Supplemental Figure 4 RNA fragment size distribution shows shorter lengths in plasma derived libraries*.

*Supplemental Figure 5 Good concordance between expected concentrations and observed TPMs*. LP = library prep replicate.

*Supplemental Figure 6 Relative RNA concentration assessed by spike-in RNA (not corrected for original biofluid input volumes)*.

*Supplemental Figure 7 t-SNE plots demonstrate the (dis)similarity of the sample biotypes*.

*Supplemental Figure 8 Log*_*10*_ *counts of the 20 most abundant genes per sample*.

*Supplemental Figure 9 Count distributions per sample*.

*Supplemental Figure 10 Gene body coverage shows typical total RNA sequencing coverage of fragmented RNA*.

*Supplemental Figure 11 Overlap of expressed genes for ePRP and ePFP*. The ePRP unique genes show an equal distribution compared to the overlapping genes, while the ePFP unique genes are lower distributed.

## Author Contributions

## References

1. Norwitz, E. R. & Levy, B. Noninvasive Prenatal Testing: The Future Is Now. Rev. Obstet. Gynecol. 6, 48–62 (2013).

2. Siravegna, G., Marsoni, S., Siena, S. & Bardelli, A. Integrating liquid biopsies into the management of cancer. Nat. Rev. Clin. Oncol. 14, 531–548 (2017).

3. Buschmann, D. et al. Toward reliable biomarker signatures in the age of liquid biopsies - how to standardize the small RNA-Seq workflow. Nucleic Acids Res. 44, 5995–6018 (2016).

4. Vo, J. N. et al. The Landscape of Circular RNA in Cancer. Cell 176, 869–881.e13 (2019).

5. Qin, Y. et al. High-throughput sequencing of human plasma RNA by using thermostable group II intron reverse transcriptases. RNA 22, 111–128 (2016).

6. Nikitina, A. S. et al. Datasets for next-generation sequencing of DNA and RNA from urine and plasma of patients with prostate cancer. Data Brief 10, 369–372 (2016).

7. Galvanin, A. et al. Diversity and heterogeneity of extracellular RNA in human plasma. Biochimie (2019). doi:10.1016/j.biochi.2019.05.011

8. Akat, K. M. et al. Detection of circulating extracellular mRNAs by modified small-RNA-sequencing analysis. JCI Insight 4, (2019).

9. Giraldez, M. D. et al. Phospho-RNA-seq: a modified small RNA-seq method that reveals circulating mRNA and lncRNA fragments as potential biomarkers in human plasma. EMBO J. 38, (2019).

10. Gaidatzis, D., Burger, L., Florescu, M. & Stadler, M. B. Analysis of intronic and exonic reads in RNA-seq data characterizes transcriptional and post-transcriptional regulation. Nat. Biotechnol. 33, 722–729 (2015).

11. Volders, P.-J. et al. LNCipedia 5: towards a reference set of human long non-coding RNAs. Nucleic Acids Res. 47, D135–D139 (2019).

12. Savelyeva, A. V. et al. Variety of RNAs in Peripheral Blood Cells, Plasma, and Plasma Fractions. BioMed Res. Int. 2017, (2017).

13. Freedman, J. E. et al. Diverse human extracellular RNAs are widely detected in human plasma. Nat. Commun. 7, (2016).

14. Hill, A. F. et al. ISEV position paper: extracellular vesicle RNA analysis and bioinformatics. J. Extracell. Vesicles 2, (2013).

15. Page, K., Shaw, J. A. & Guttery, D. S. The liquid biopsy: towards standardisation in preparation for prime time. Lancet Oncol. 20, 758–760 (2019).

16. Hardwick, S. A. et al. Spliced synthetic genes as internal controls in RNA sequencing experiments. Nat. Methods 13, 792–798 (2016).

17. Tulkens, J. et al. Increased levels of systemic LPS-positive bacterial extracellular vesicles in patients with intestinal barrier dysfunction. Gut gutjnl-2018-317726 (2018). doi:10.1136/gutjnl-2018-31772618.

18. Van Deun, J. et al. The impact of disparate isolation methods for extracellular vesicles on downstream RNA profiling. J. Extracell. Vesicles 3, (2014).

19. Vergauwen, G. et al. Confounding factors of ultrafiltration and protein analysis in extracellular vesicle research. Sci. Rep. 7, 2704 (2017).

20. Hendrix, A. et al. Effect of the Secretory Small GTPase Rab27B on Breast Cancer Growth, Invasion, and Metastasis. JNCI J. Natl. Cancer Inst. 102, 866–880 (2010).

21. Ev-Track Consortium et al. EV-TRACK: transparent reporting and centralizing knowledge in extracellular vesicle research. Nat. Methods 14, 228–232 (2017).

22. Martin, M. Cutadapt removes adapter sequences from high-throughput sequencing reads. EMBnet.journal 17, 10 (2011).

23. Dobin, A. et al. STAR: Ultrafast universal RNA-seq aligner. Bioinformatics 29, 15–21 (2013).

24. Li, H. et al. The Sequence Alignment/Map format and SAMtools. Bioinformatics 25, 2078–2079 (2009).

25. Zerbino, D. R. et al. Ensembl 2018. Nucleic Acids Res. 46, D754–D761 (2018).

26. Bray, N. L., Pimentel, H., Melsted, P. & Pachter, L. Near-optimal probabilistic RNA-seq quantification. Nat. Biotechnol. 34, 525–527 (2016).

27. Love, M. I., Huber, W. & Anders, S. Moderated estimation of fold change and dispersion for RNA-seq data with DESeq2. Genome Biol. 15, 550 (2014).

28. Zhang, X. O. et al. Diverse alternative back-splicing and alternative splicing landscape of circular RNAs. Genome Res. 26, 1277–1287 (2016).

