## Supplemental Figure for "Performance assessment of total RNA sequencing of human biofluids and extracellular vesicles"

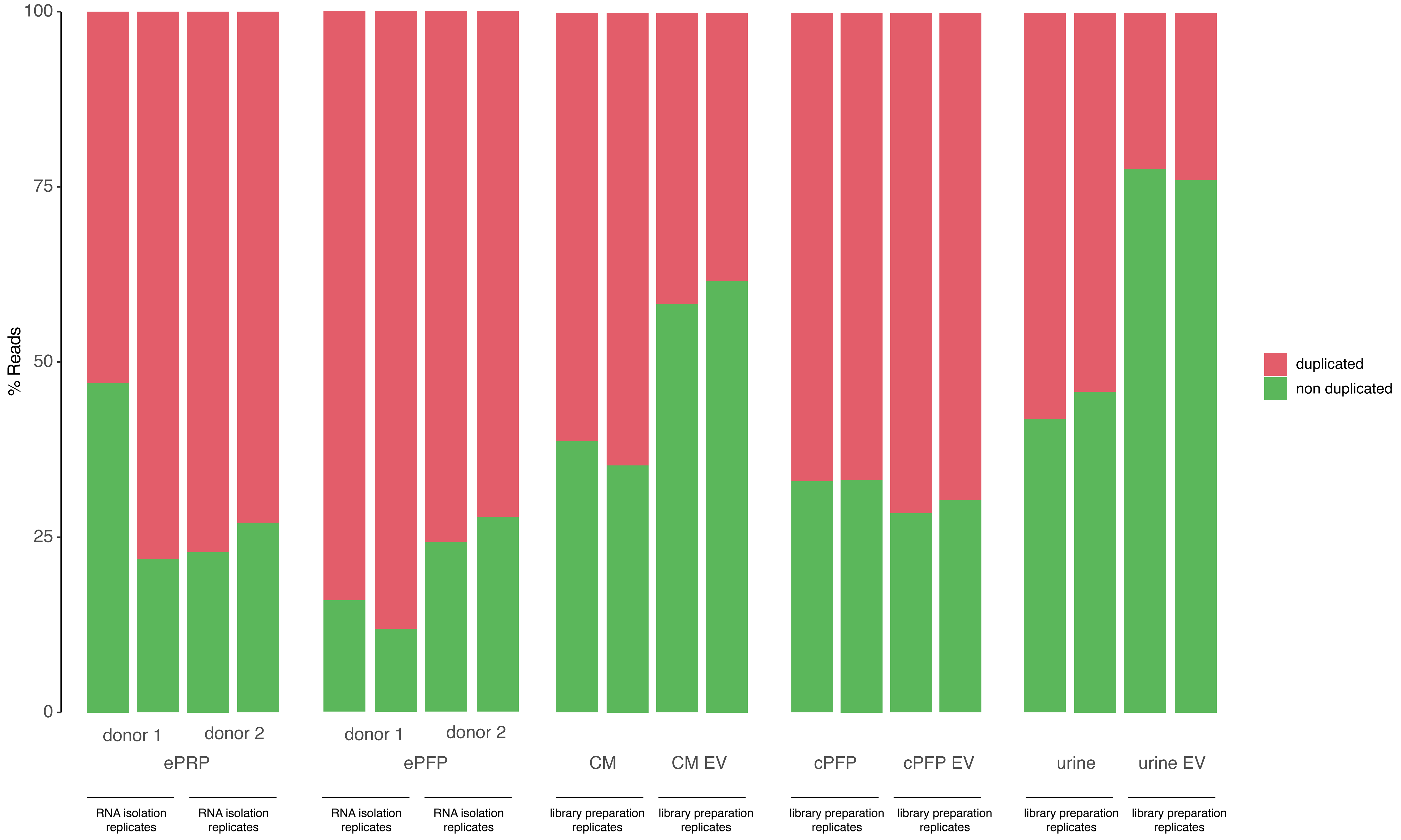


**Supplemental Figure 1 Read duplication levels are markedly different among different biomaterials.**

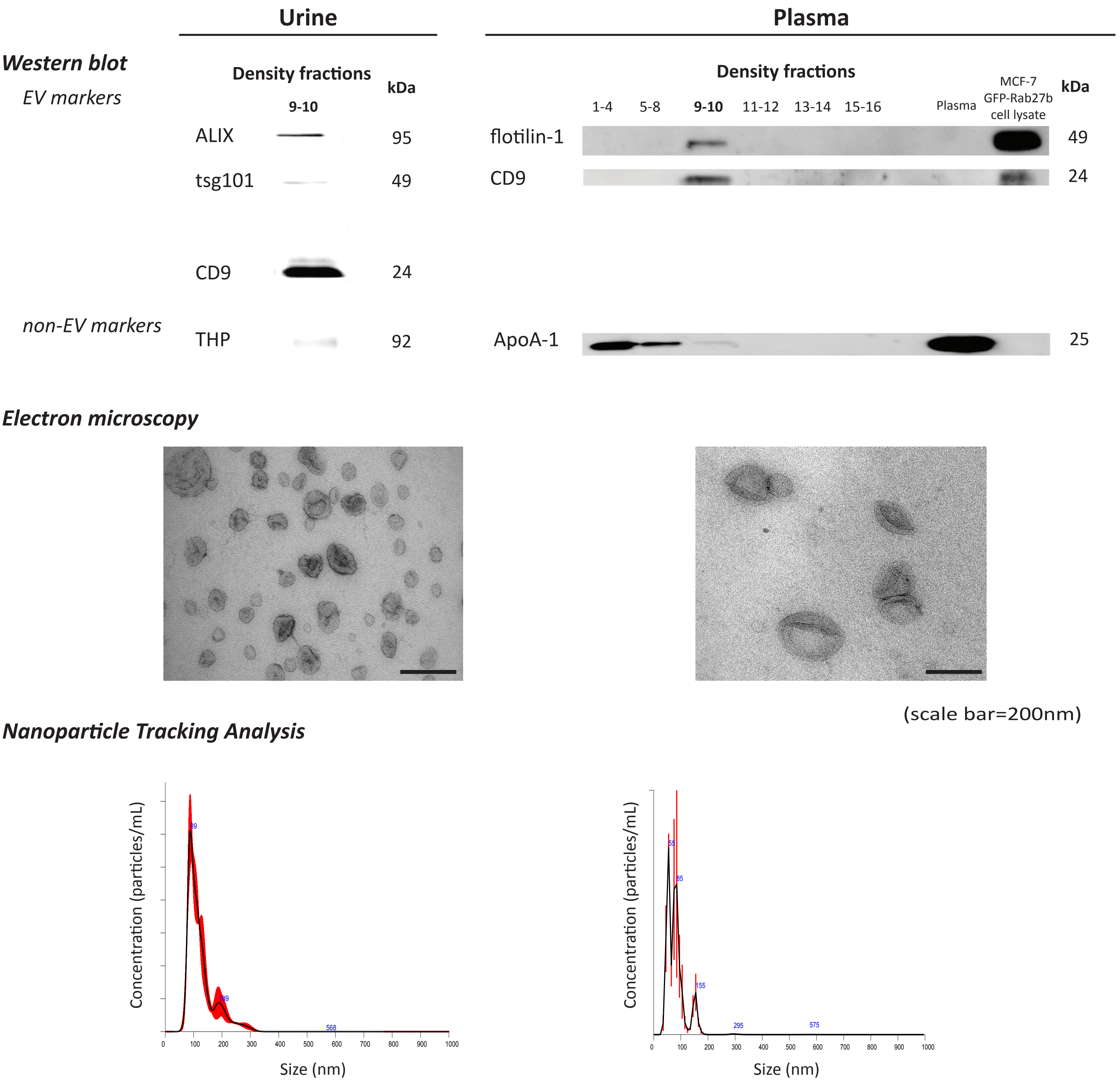


**Supplemental Figure 2a Characterization of EV from urine and plasma samples.** Proteins are analyzed by western blot using specific EV markers (ALIX, tsg101, CD9 and flotillin-1) and non-EV markers (THP and ApoA-1). EV samples (density gradient fractions 9-10) are enriched in EV proteins and depleted for contaminants. EVs were qualitatively and quantitatively analyzed by electron microscopy and nanoparticle tracking analysis.

### MCF-7 GFP-Rab27b

#### *Western blot*

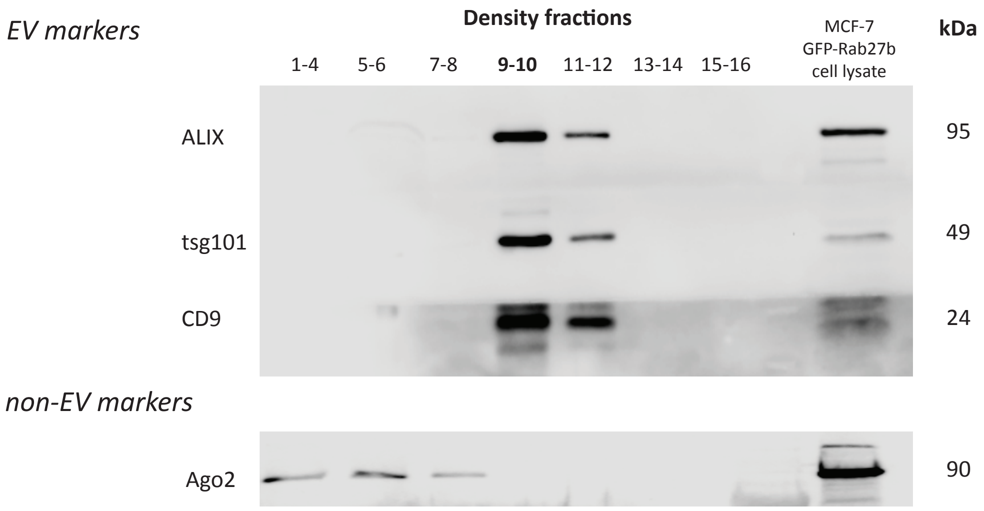


#### *Electron microscopy*

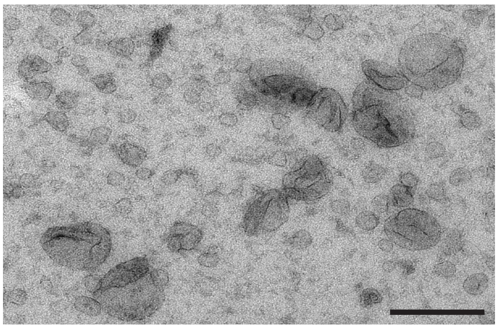

(scale bar=200nm)

#### *Nanoparticle Tracking Analysis*

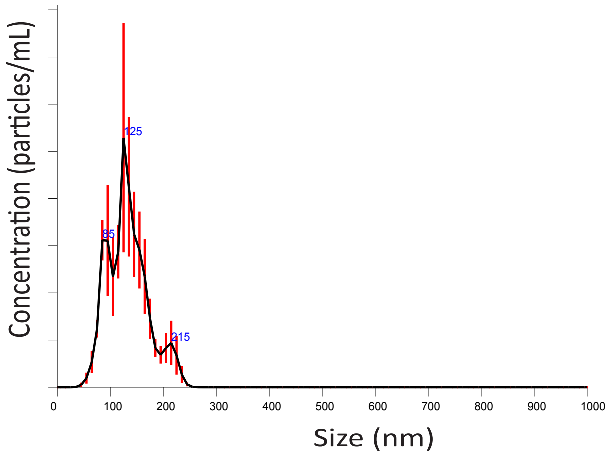


**Supplemental Figure 2b Characterization of EV from MCF-7 GFP-Rab27b cells.** Proteins are analyzed by performing western blot using specific EV markers (ALIX, tsg101 and CD9) and non-EV markers (Ago2). EV samples (density gradient fractions 9-10) are enriched in EV proteins and depleted for contaminants. EVs were qualitatively and quantitatively analyzed by electron microscopy and nanoparticle tracking analysis.

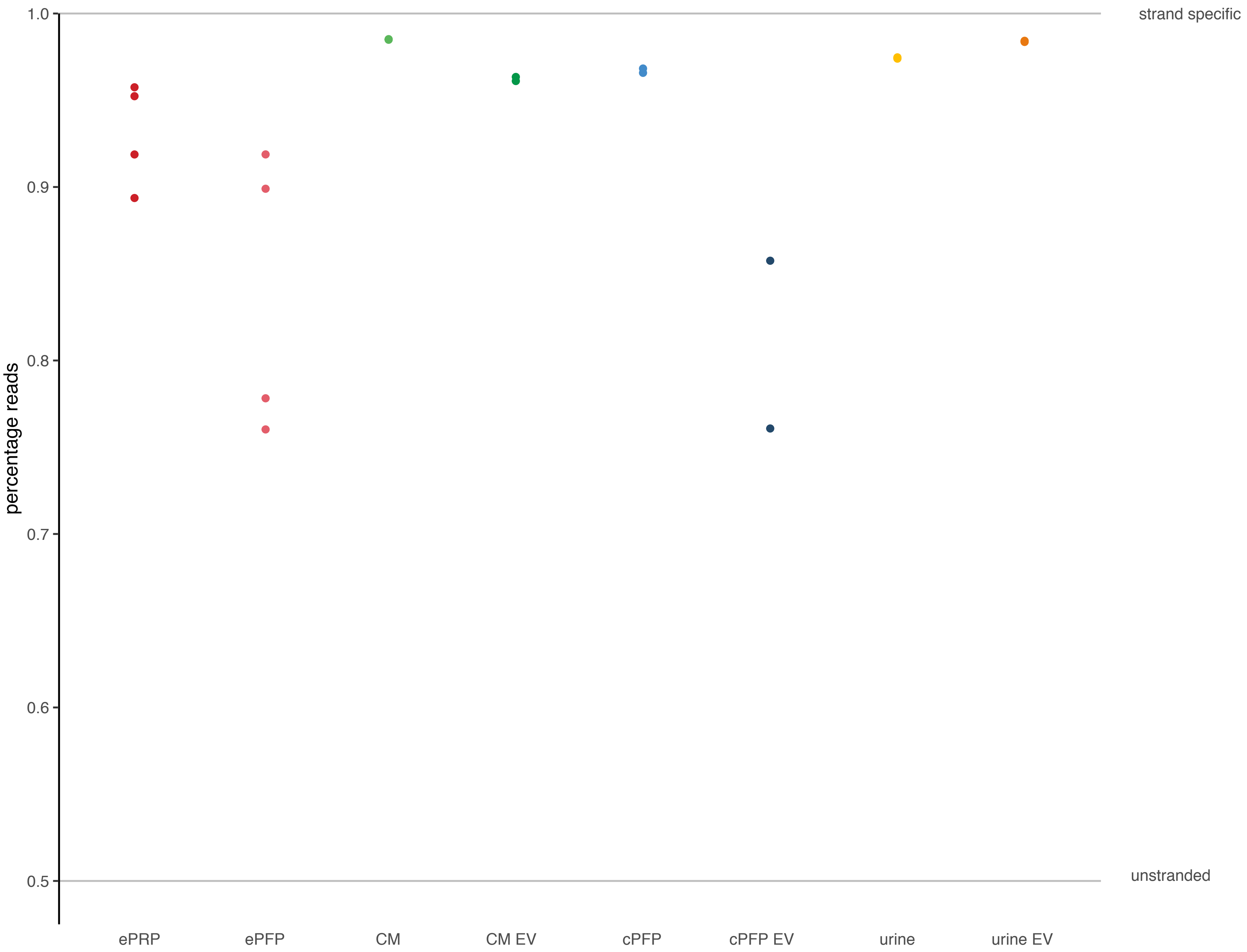


Supplemental Figure 3. Percentage of reads originating from the sense strand to demonstrate good strandedness of the kit.

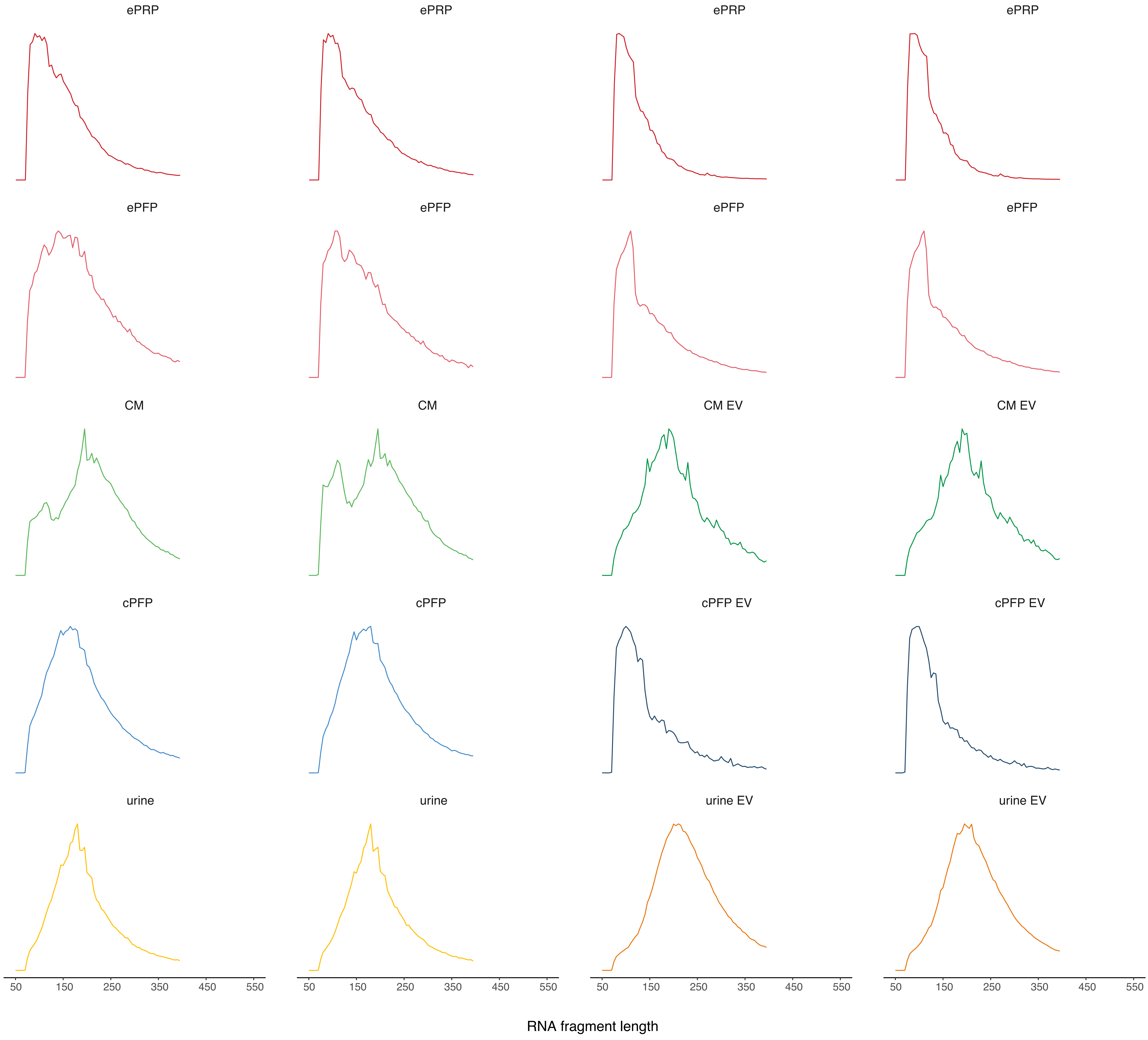


Supplemental Figure 4 RNA fragment size distribution shows shorter lengths in plasma derived libraries.

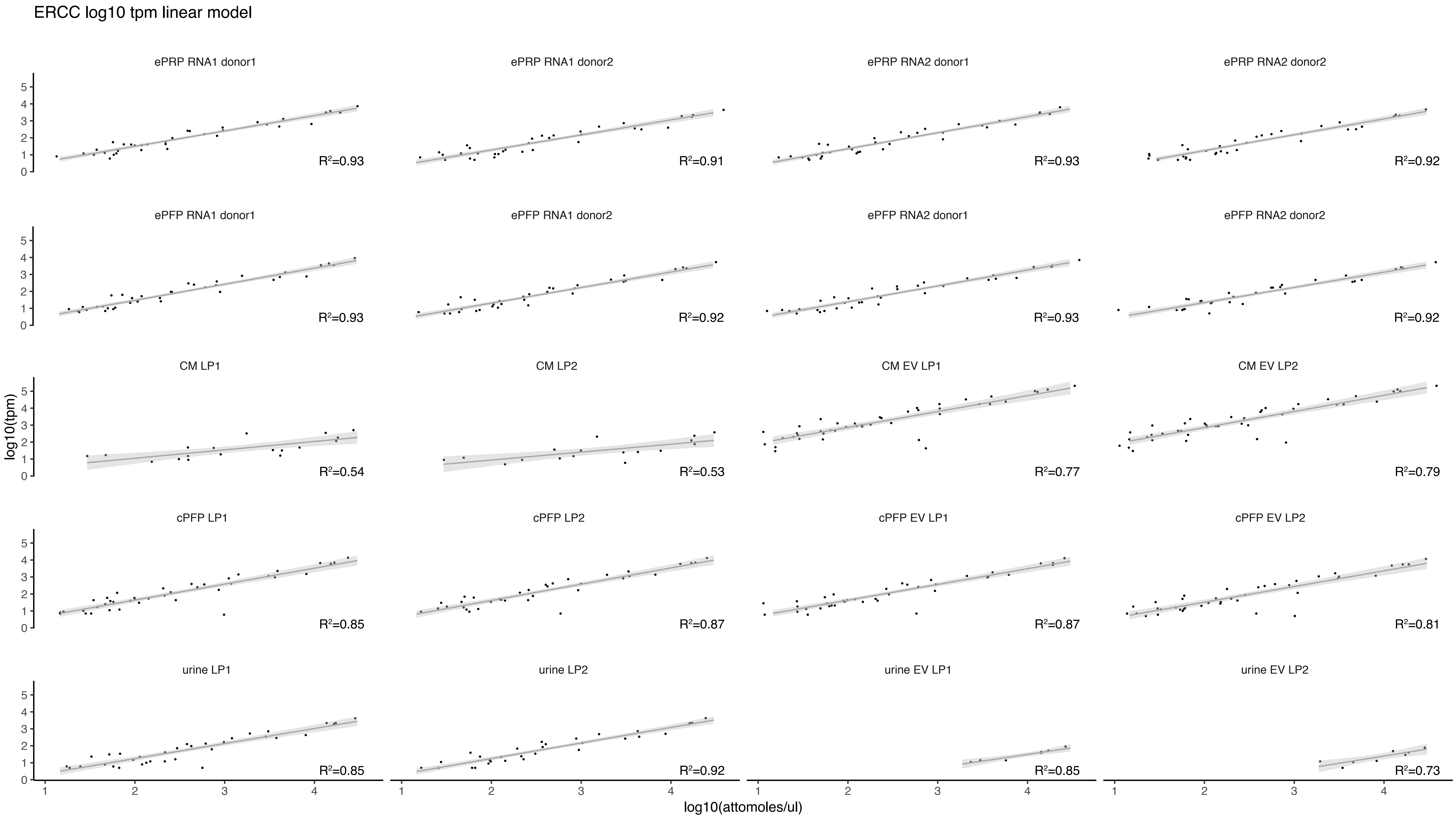


Supplemental Figure 5 Good concordance between expected concentrations and observed TPMs. LP = library prep replicate.

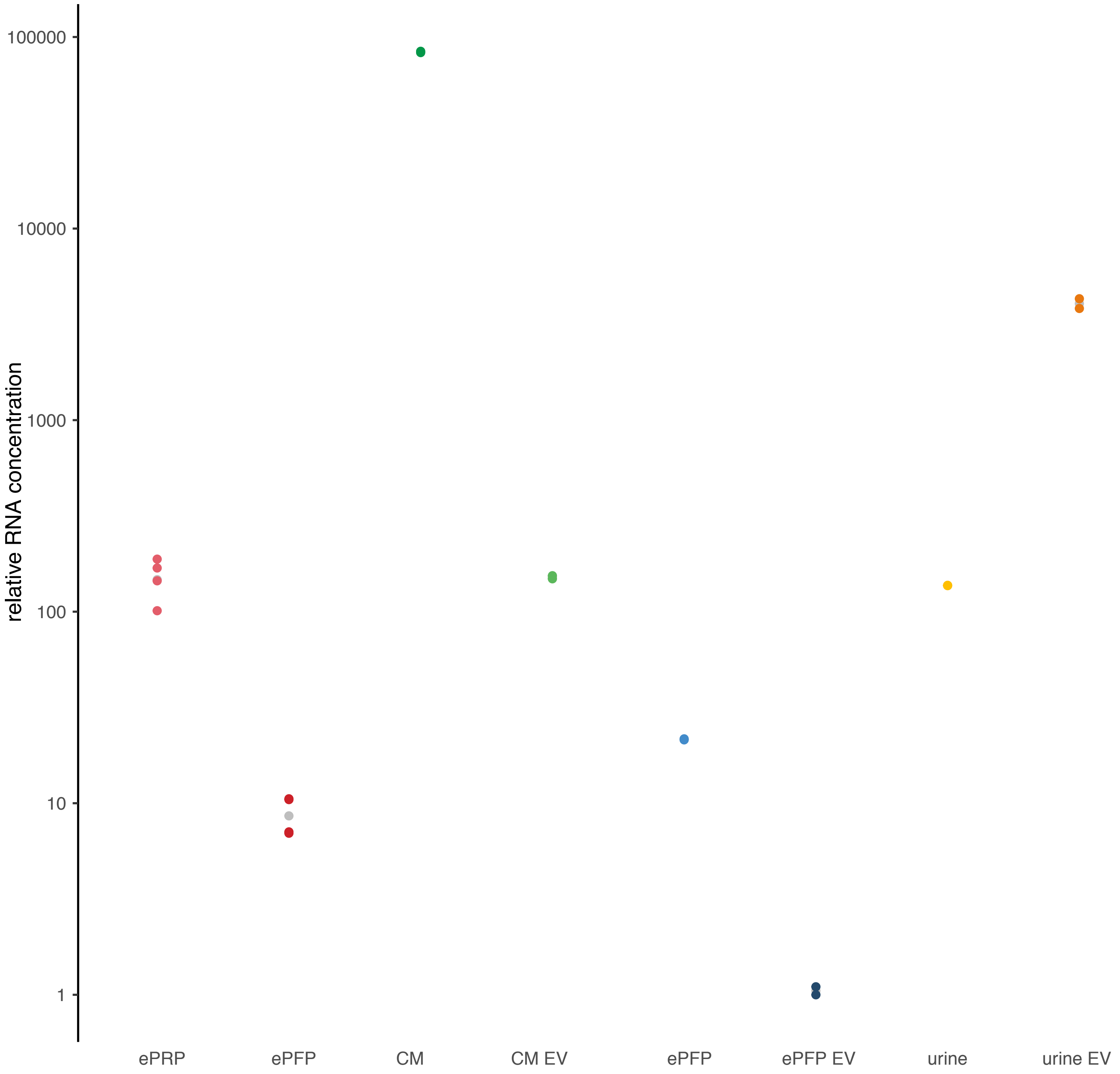


Supplemental Figure 6 Relative RNA concentration assessed by spike-in RNA (not corrected for original biofluid input volumes).

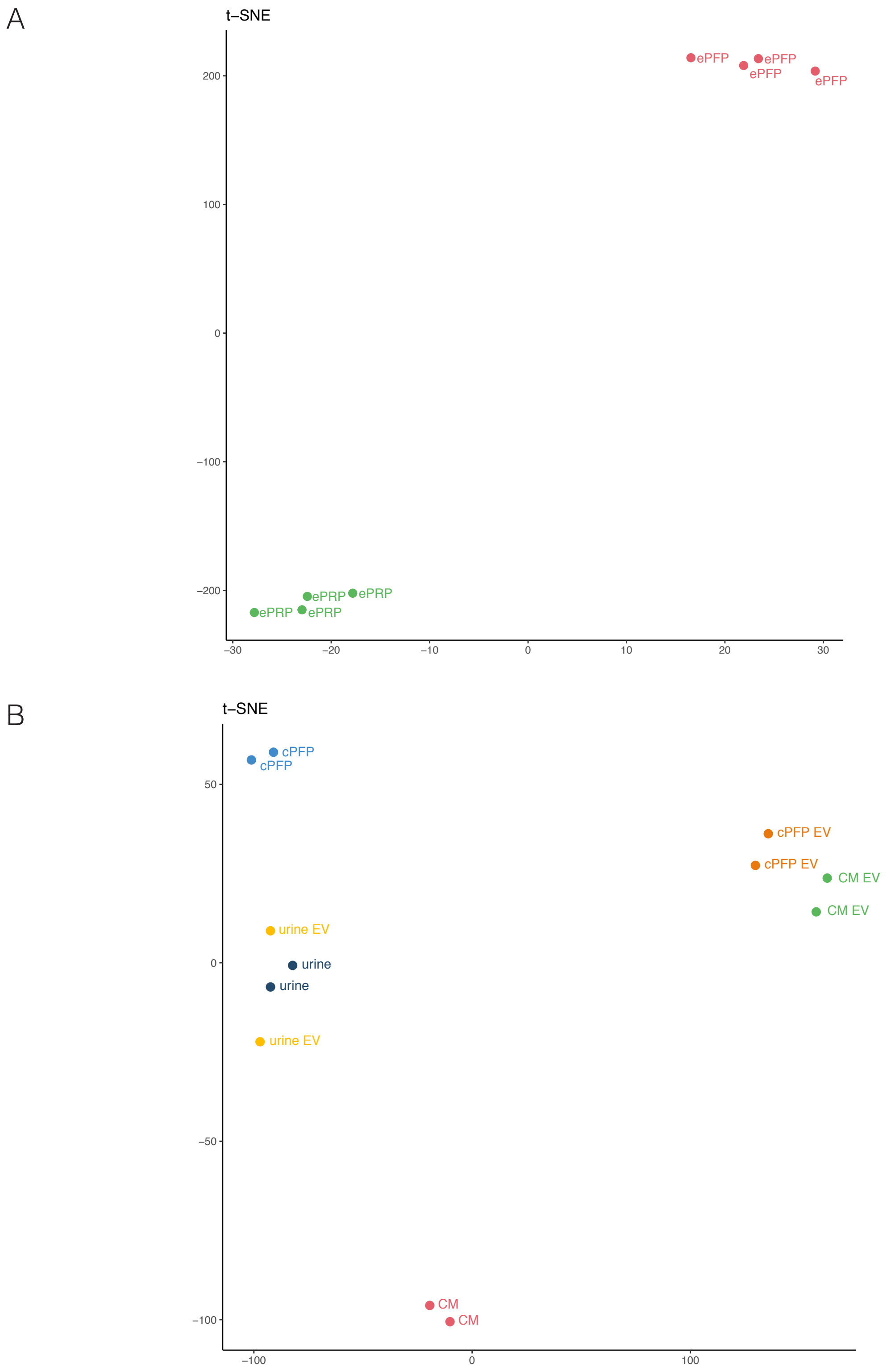


Supplemental Figure 7 t-SNE plots demonstrate the (dis)similarity of the sample biotypes.

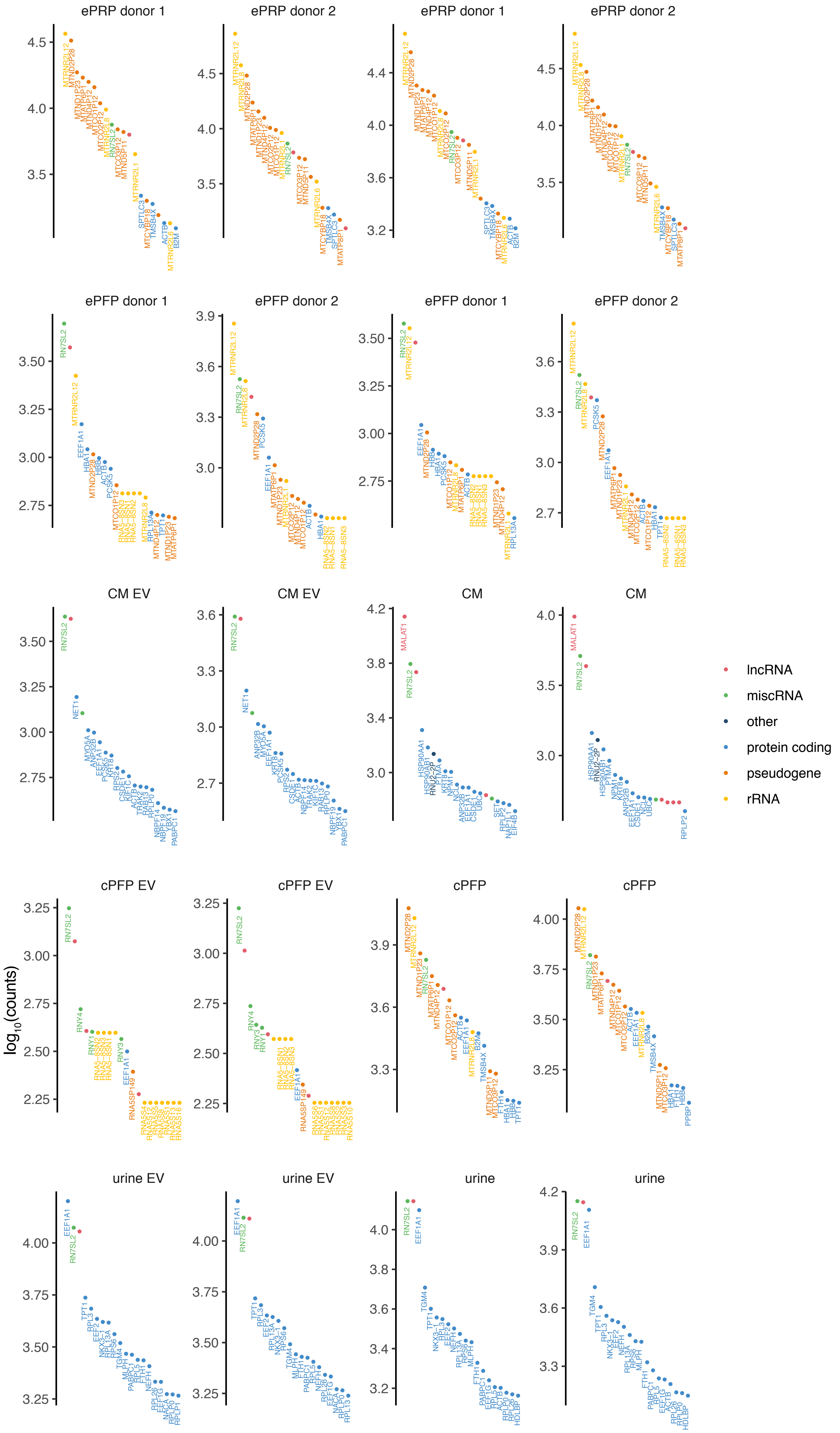


Supplemental Figure 8 $\text{Log}_{10}$ counts of the 20 most abundant genes per sample.

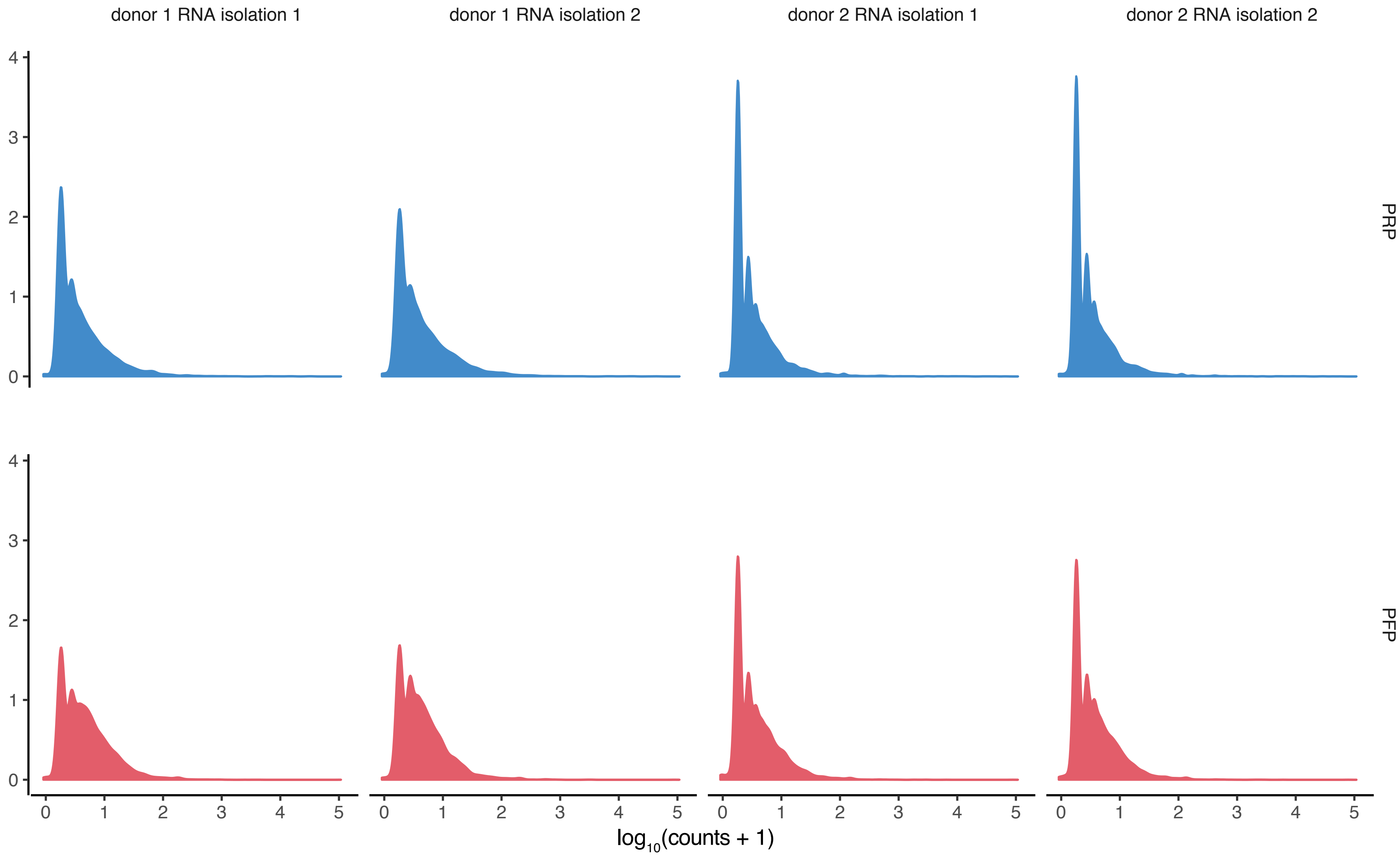


Supplemental Figure 9 Count distributions per sample.

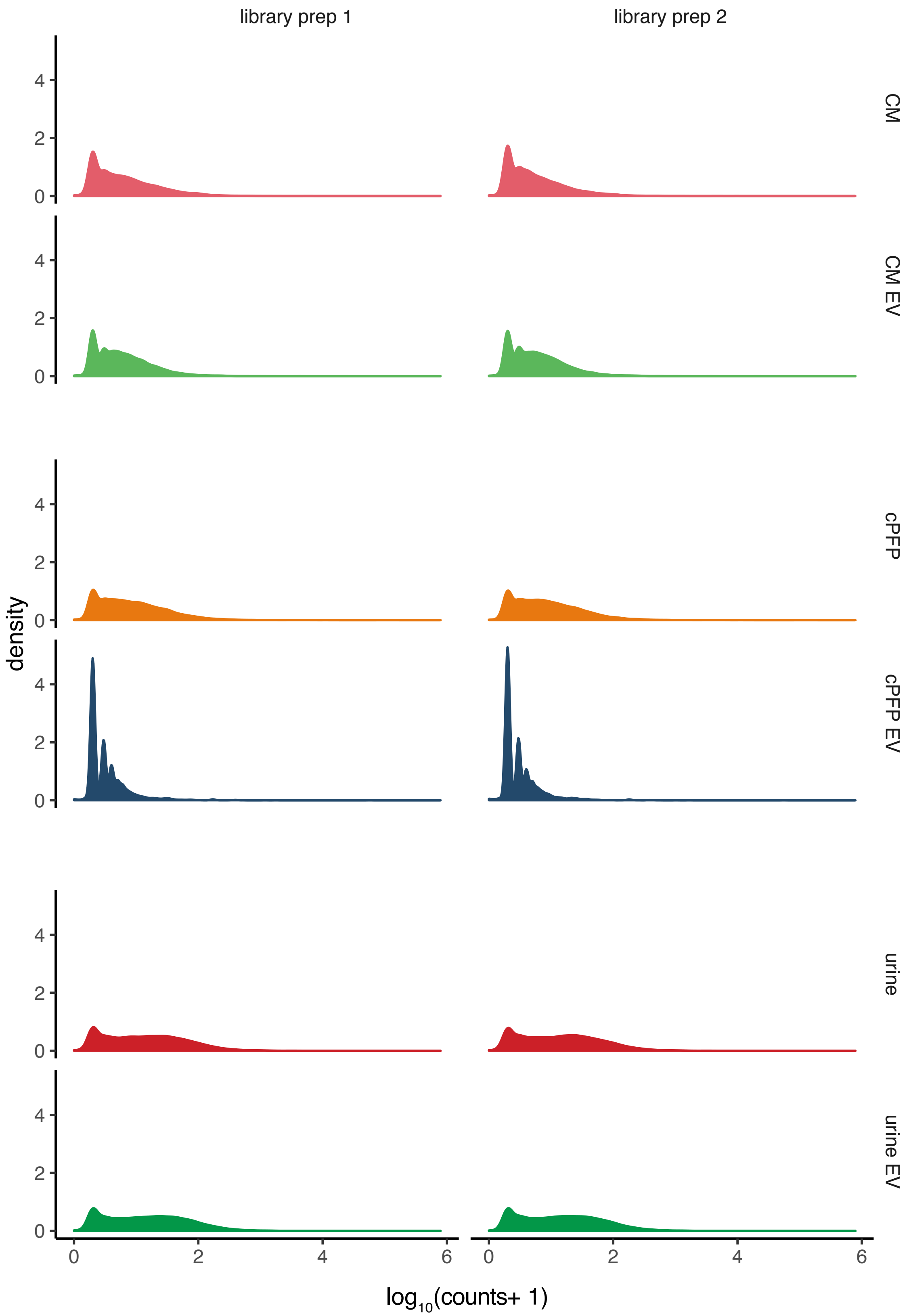


Supplemental Figure 9 Count distributions per sample.

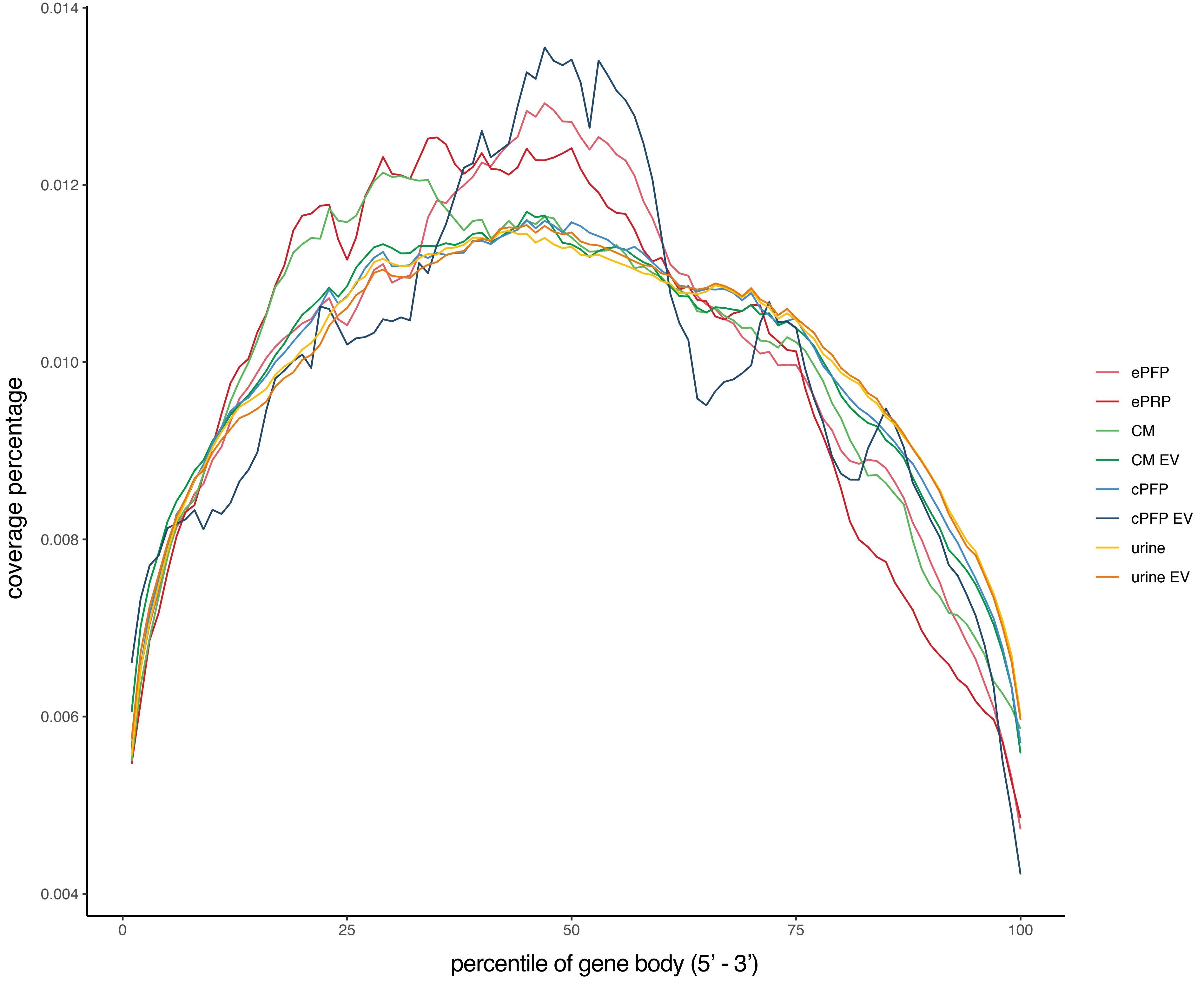


Supplemental Figure 10 Gene body coverage shows typical total RNA sequencing coverage of fragmented RNA.

A

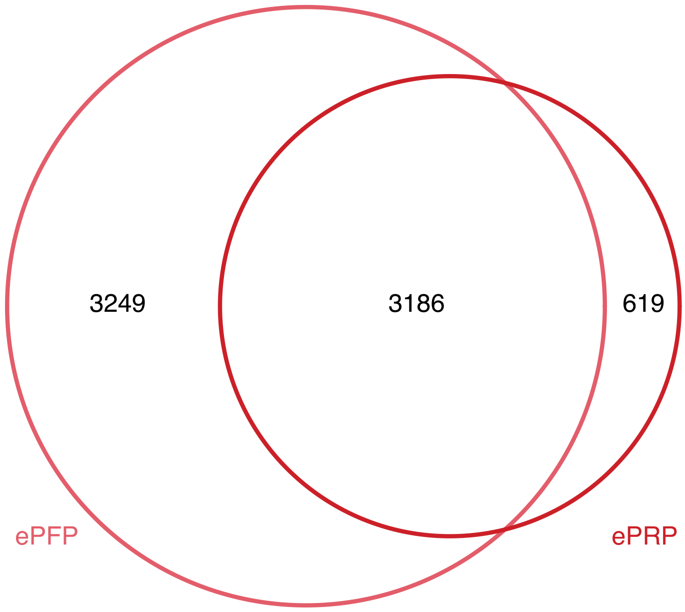


B

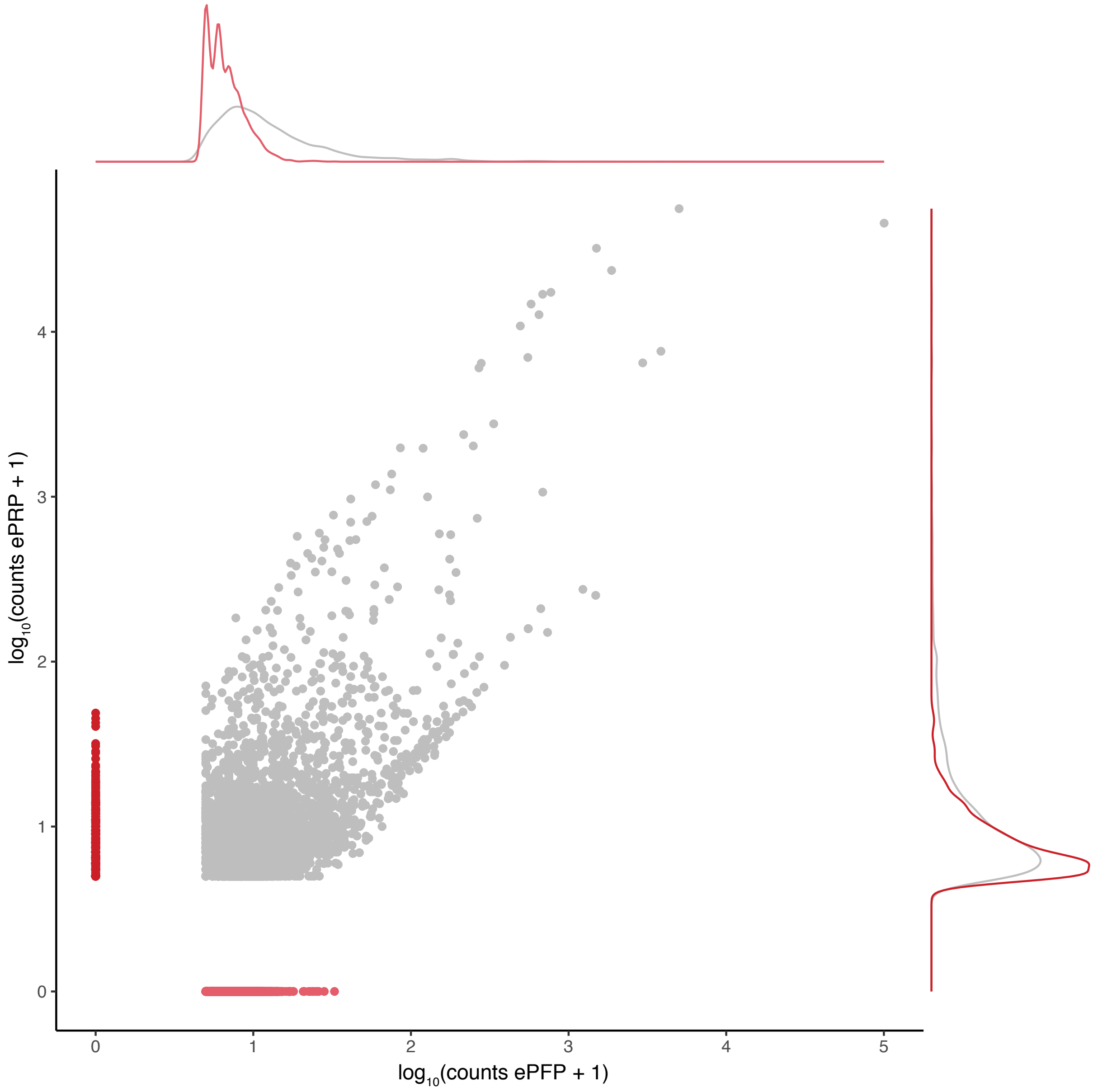


**Supplemental Figure 11 Overlap of expressed genes for ePRP and ePFP.** The ePRP unique genes show an equal distribution compared to the overlapping genes, while the ePFP unique genes are lower distributed.
